# The Great Migration and African-American genomic diversity

**DOI:** 10.1101/029173

**Authors:** Soheil Baharian, Maxime Barakatt, Christopher R. Gignoux, Suyash Shringarpure, Jacob Errington, William J. Blot, Carlos D. Bustamante, Eimear E. Kenny, Scott M. Williams, Melinda C. Aldrich, Simon Gravel

## Abstract

Genetic studies of African-Americans identify functional variants, elucidate historical and genealogical mysteries, and reveal basic biology. However, African-Americans have been under-represented in genetic studies, and little is known about nation-wide patterns of genomic diversity in the population. Here, we present a comprehensive assessment of African-American genomic diversity using genotype data from nationally and regionally representative cohorts. We find higher African ancestry in southern United States compared to the North and West. We show that relatedness patterns track north- and west-bound routes followed during the Great Migration, suggesting that admixture occurred predominantly in the South prior to the Civil War and that ancestry-biased migration is responsible for regional differences in ancestry. Rare genetic traits among African-Americans can therefore be shared over long geographic distances along the Great Migration routes, yet their distribution over short distances remains highly structured. This study clarifies the role of recent demography in shaping African-American genomic diversity.

## Introduction

The history of African-American populations is marked by dramatic migrations within Africa, through the transatlantic slave trade, and within the United States (US). By 1808, when the transatlantic slave trade was made illegal in the US, approximately 360,000 Africans had been brought forcibly into the US in documented voyages (*1*). International and domestic slave trade continued to impose long-distance migration on enslaved African-Americans until the end of the Civil War, in 1865. By 1870, the US census reported 4.88 million “colored” individuals of which 90% lived in the South (*2*).

Despite the ban on slavery, economic and social perspectives for most African-Americans remained bleak. Better opportunities in the North (Northeast and Midwest) and West led millions of African-Americans to leave the South between 1910 and 1970 (*3*). This demographic event known as the Great Migration profoundly reshaped African-American communities across the US (4). Today, 45 million Americans identify as Black or African-American.

A history of slavery and of systemic discrimination led to increased social, economic, and health burdens in many African-American communities. Health disparities continue to be compounded by poverty, unequal access to care, and unequal representation in medical research. To reduce health disparity in research, many cohorts are currently being assembled to encompass more of the diversity within the US (*5, 6*). These cohorts create opportunities in both medical and population genetics; they also require an understanding of genetic diversity within diverse cohorts. However, the large-scale migrations and incomplete genealogical records for African-Americans present a challenge for such an understanding. Previous studies have described the proportions of African, European, and Native American ancestries across individuals (*7-13*), the amount of diversity in sequence data (*9, 14, 15*) and inferred admixture models (*12, 16, 17*). However, the genetic population structure among African-Americans is not well understood.

Here, we use cohorts including 3,726 African-Americans and a total of 13,199 individuals geographically distributed across the contiguous US to investigate nation-wide population structure among African-Americans. We first confirm and refine previous estimates of admixture proportions and timing in the population, and find significant differences in ancestry proportions between US regions. We then investigate relatedness among African-Americans and European-Americans through identity-by-descent analysis, and identify long- and short-range patterns of isolation-by-distance. We introduce quantitative models, incorporating both census data and fine-scale migration, to describe these isolation-by-distance patterns and infer migratory patterns in the population. Integrating quantitative models for admixture, relatedness information, and historical data, we identify ancestry-biased migrations during the Great Migration as a driving force for ancestry and relatedness variation among African-Americans. The analysis of geographically distributed cohorts through detailed mathematical modeling therefore helps us understand the distribution of genetic diversity in large cohorts and provides new insights into recent human demography.

## Cohorts

We analyzed data from three cohorts: (a) Health and Retirement Study (HRS), with 1,501 African-Americans and 9,308 European-Americans sampled representatively across all US states, and including urban and rural regions; (b) Southern Community Cohort Study (SCCS), including 2,128 African-Americans sampled within the southern US in rural locations; (c) 1000 Genomes Project cohort of 97 individuals of African ancestry from the southwest USA (ASW). (For detailed information, see Methods and Tables S4 and S5.)

## Admixture patterns

Individual genomes carry genetic material from multiple ancestral lineages, and each diploid locus derives ancestry from two distinct lineages. We used RFMix (*11*) together with 1000 Genomes panels from Africa, Europe, and Asia to identify the most likely continental location of the pre-1492 ancestors that contributed genetic material at each locus for individuals in the cohorts (Fig. 1D, Fig. S14, and Methods). The overall proportion of African ancestry (Table 1) is substantially higher in the SCCS and HRS than in the ASW and the recently published 23andMe cohort (*12*).

**Table 1:** Inferred proportions of African, European, and Native American/Asian Ancestry in three African-American cohorts.

| Cohort | % African | % European | % Native American |
| --- | --- | --- | --- |
| SCCS | 84.1 | 14.7 | 1.1 |
| HRS | 82.1 | 16.7 | 1.2 |
| ASW | 75.9 | 21.3 | 2.8 |

**Figure 1:**
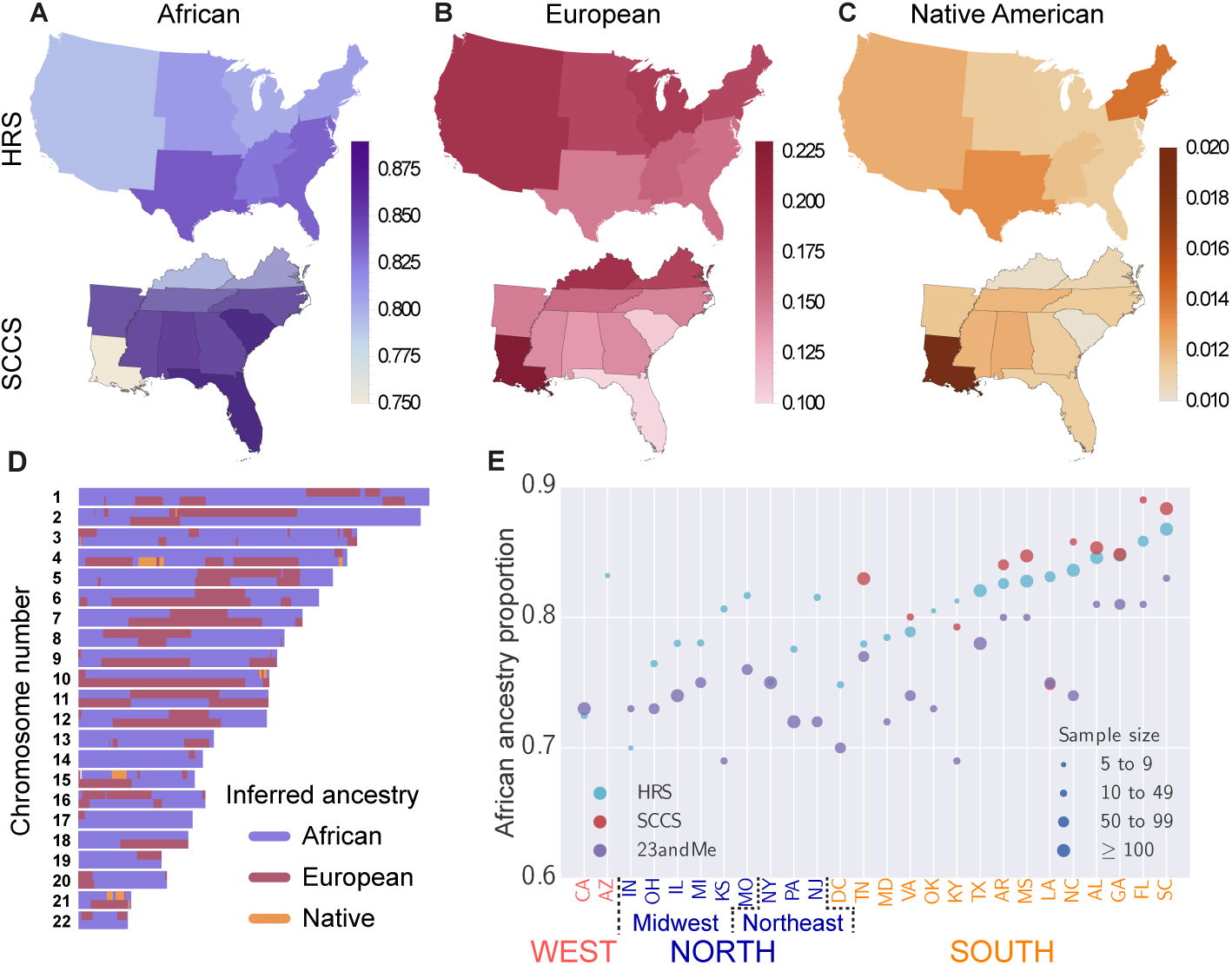
Inferred regional ancestry proportions for the HRS and SCCS cohorts: (A) African, (B) European, and (C) Native American ancestries. (D) Local ancestry assignment along the autosomes for an African-American individual from HRS. (E) Comparison of the African ancestry proportions in the HRS, SCCS, and 23andMe stratified by state. 23andMe proportions are from Ref. (*12*) and are reported for ease of comparison.

The HRS cohort can be thought of as representative of the entire African-American population, while the SCCS focuses primarily on individuals attending community health centers in rural, underserved locations in the South. The sampling for the ASW and 23andMe did not aim for specific representativeness (see Methods). In the HRS, average African ancestry proportion is 83% in the South and lower in the North (80%, bootstrap *p* = 6 × 10^−6^) and West (79%, *p* = 10^−4^) (Fig. 1). Within the SCCS, African ancestry proportion is highest in Florida (89%) and South Carolina (88%) and lowest in Louisiana (75%) with all three significantly different from the mean (Florida *p* = 0.006, South Carolina *p* = 4 × 10^−4^, and Louisiana *p* < 10^−5^; bootstrap). The elevated African ancestry proportion in Florida and South Carolina is also observed in the HRS and in the 23andMe study (*12*), but Louisiana is more variable across cohorts (Fig. 1E). As expected, European ancestry proportions largely complement those of African ancestry across the US.

Recombination breaks down ancestral haplotypes over time (Fig. 1D). We inferred the timing of admixture based on the length of continuous ancestry segments along individual genomes using Tracts (*16*). Since there are few real Native American segments, even a small number of spurious Native American segments can bias the inference. Thus, we first considered a model with two source populations: African and non-African. Assuming a single admixture event, we estimated the time of admixture onset *g*, where *g* = 1 means that the parents of the individual are the founders of the admixed population and that the current individual represents the first admixed generation. For HRS, we inferred a timing of *g* = 5.8 generations ago (Fig. S9; confidence interval (CI) in Table S3). The estimated year of birth of the first admixed children is *T* = *T_s_* − (*g* − 1)*τ*, where *T_s_* = 1939.8 is the average year of birth of HRS individuals and *τ* is the generation time. Individuals born *τ* years earlier should be 1 generation closer to the onset of admixture. Correlating birth year and inferred admixture time within our cohort (Fig. 2D), we inferred *τ* = 27.4 (*r*^2^ = 0.88, *p* = 10^−7^), which leads to an admixture year of 1808.

A model allowing for two phases of European admixture outperforms the single-pulse model for HRS and SCCS (see Methods). In HRS, it suggests a first admixture event in 1740 (8.3 generations ago) and a second pulse, of approximately equal size, in 1863 (3.8 generations ago) (Fig. S9 and Methods). Mean birth year in SCCS is *T_s_* = 1946.9, supporting a single admixture event in 1802 (6.3 generations ago), or two events in 1714 and 1854 (9.5 and 4.4 generations ago) (Fig. 2A, Fig. S9, and Methods). The two-pulse model is a coarse simplification of the historical admixture process, but the data strongly supports ongoing admixture, predominantly before or around the end of the Civil War. The limited role of early 20th century admixture is further supported by the similarity in the inferred single-pulse time to admixture in all HRS census regions (between 5.4 and 6.2 generations ago, Fig. S18) and all cohorts, which is easily explained if most admixture occurred in the South prior to the Great Migration. The similar levels of African ancestry for all age cohorts also supports limited European admixture between 1930 and 1960 (Fig. 2D). Importantly, more recent admixture is not represented in the SCCS and HRS cohorts; only two participants were born after 1970.

**Figure 2:**
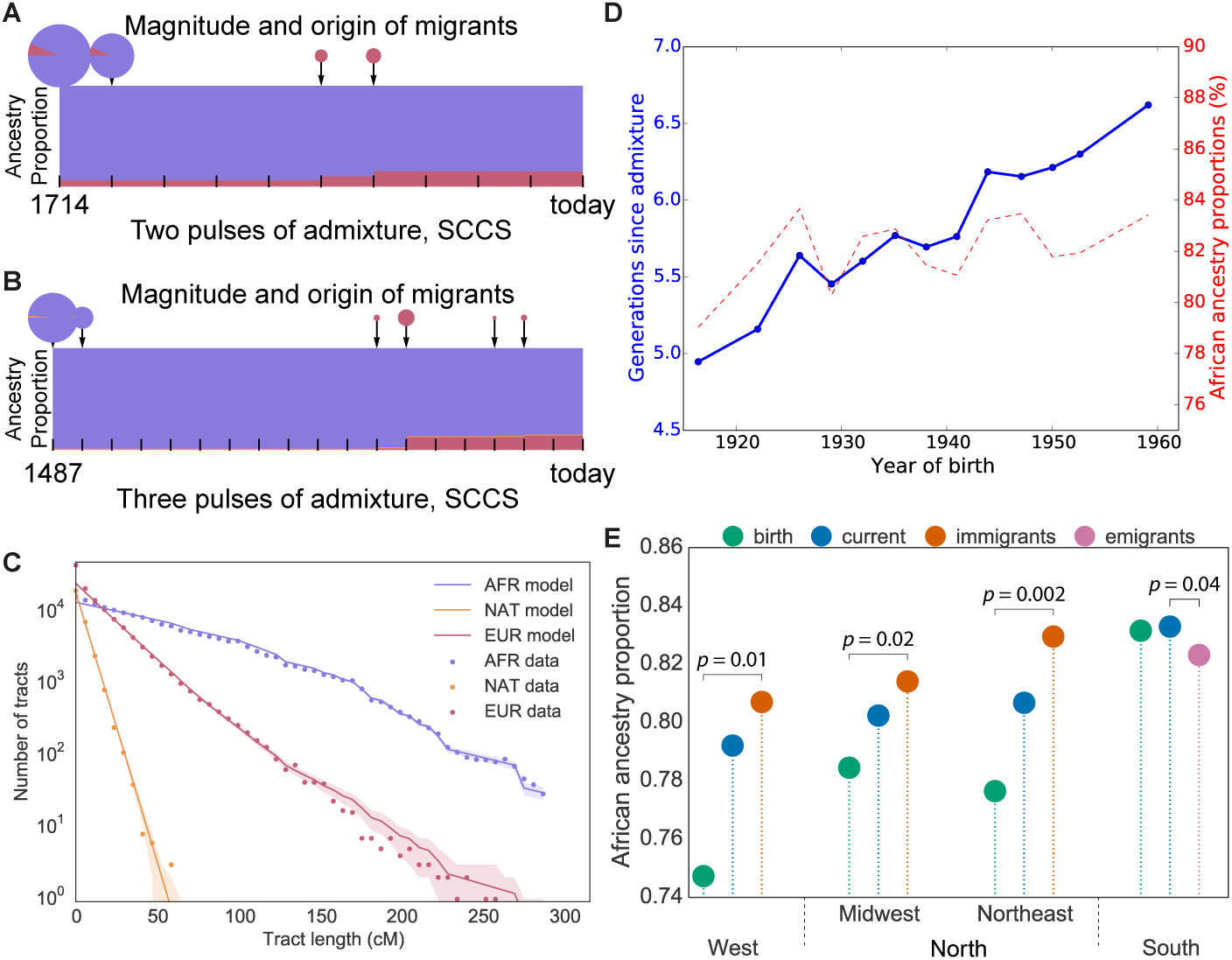
Admixture times and proportions of ancestral populations for SCCS in (A) the model with two pulses of admixture and (B) the model with three pulses of admixture. (C) Distribution of continuous ancestry tract lengths (dots) compared with predictions from the best-fit model (lines) Points in the shaded area are within one standard deviation of the predicted result. (D) Inferred time to admixture and African ancestry proportions as functions of birth year in HRS African-Americans. (E) Proportions of African ancestry in African-Americans within the North, South, and West using region of birth, region of residence, and migration status; bootstrap p-values are calculated between disjoint sets of individuals.

Time estimates point to admixture occurring when most ancestors to present-day African-Americans lived in the South. Regional differences in ancestry are therefore unlikely to be caused by differences in recent admixture rates, and the large influx of migrants from the South would have strongly attenuated any earlier differences. An alternate explanation for regional differences in ancestry proportions is ancestry-biased migrations, if individuals with higher European ancestry were more likely to migrate to the North and West during the Great Migration. To validate the ancestry-biased migration model, we compared ancestry proportions of HRS individuals according to their region of birth, residence, and migration status.

European ancestry proportions in African-americans who left the South (16.5%) is elevated compared to individuals who remained in the South (15. 3%, bootstrap *p* = 0.04), confirming that ancestry-biased migrations continued at least to the mid-20th century. These migrants had substantially *less* European ancestry than people already established in the North (20.9%) and West (25.0%) (Fig. 2E). Since the latter two groups received large contributions from the first generation of migrants, the excess European ancestry suggests a stronger ancestry bias in the first wave of migration. This change over time in ancestry-biased migration is consistent with historical accounts that southern African-American migrants to northern cities during the later stages of the Great Migration had darker complexion than North-born African-Americans (see (*18*), p. 179). The change could be explained by better social opportunities available to individuals with higher levels of European ancestry: Individuals with wealth and education were much more likely to migrate in the first wave of the migration (see (*18*), p. 167). Despite the ongoing ancestry bias, the migrations of HRS participants led to more uniform ancestry proportions across regions. Interestingly, the proportion of African ancestry among African-Americans *increased* in all four US regions; the ancestry bias caused migrants to have levels of admixture between those of the South-born and North-born individuals. Their departures and arrivals both increased the regional African ancestry proportions.

Out of 1,491 non-Hispanic African-Americans in HRS, 11 individuals have more than 5% Native American ancestry. Within SCCS, this proportion is only 8 out of 2,128 individuals. The ASW cohort, with 8 out of 97 individuals above this threshold, is a clear outlier; however, the other 89 individuals have similar amounts of Native American ancestry to the other studies: If we filter out individuals contributing more than 5% Native American ancestry from each cohort, the proportion of Native American ancestry in the remaining individuals is close to 1.1% in the SCCS, in all HRS census regions, and in the ASW. The filtered SCCS Louisianans have significantly more Native American ancestry (1.6%, bootstrap *p* = 2 × 10^−5^), and South Carolinians have less (0.09%, *p* = 2 × 10^−5^). We did not find a global correlation between European and Native American ancestry, except within Louisiana (Fig. S7).

A three-population admixture model accounting for Native American admixture confirmed the continuous, relatively early European admixture, and suggested that Native American admixture occurred earlier, consistent with previous findings of (*12*). Inferred dates of admixture are 1487 for the SCCS (Figs. 2B and 2C) and 1496 for the HRS (Figs. S10 and S11), as described in Methods. We suspect presence of a small amount of spurious, short segments of inferred Native American ancestry could bias the inference toward these unrealistically early dates. The lack of longer Native American segments nevertheless suggests that most Native American ancestry in African-Americans results from contact in the early days of slavery (see, e.g., (*19*)).

Along the X chromosome in the HRS, we estimate 84.82% African ancestry, 12.89% European ancestry, and 2.29% Native American ancestry (bootstrap 95% CI [2.14%, 2.45%]). The higher proportion of African ancestry along the X compared to autosomes is consistent with previous studies and the historical records of admixture occurring predominantly through European-American males admixing with African-American females (*17*). A model with a single pulse of admixture (as considered in (12)) applied to the present data suggests 28.6% Europeans among male contributors, but only 5.2% among female contributors. By contrast, it suggests almost no contribution from Native American males, and 3% from Native American females.

The US Census includes a separate category for Hispanic/non-Hispanic ethnicity. In HRS, 32 African-Americans have self-identified as Hispanics (of which only 10 are within the contiguous US). Genetic ancestry within this group is distinct from the bulk of the non-Hispanic African-American population in at least two ways: elevated Native American ancestry and a higher genetic similarity to southern European populations (Figs. S16 and S17). The correlation between southern European and Native American ancestries also holds in individuals who do not self-identify as Hispanic, particularly in Louisiana (see Methods). Individuals with elevated Native American and southern European ancestry would not be identified by self-reported ethnicity or by genetic estimates of African/non-African ancestry, yet they may have distinct response patterns to medical tests (*20, 21*).

## Identity by descent

The classical isolation-by-distance model predicts that genetic relatedness between individuals decreases as their geographic distance increases (*22*). However, large-scale migrations can dramatically alter this picture (*23*). To investigate the effect of recent migrations on patterns of genetic relatedness within African-Americans, we consider genetic segments that are identical-by-descent (IBD) between pairs of individuals. Long IBD segments (*l* ≥ 18 cM) correspond to an expected common ancestor living within the last 8 generations and are informative of very recent demography (see Methods).

Figures 3A-B and S20-S23 show the mean pairwise relatedness among seven geographic regions in the US for African-Americans and European-Americans. Here, the relatedness of two individuals is defined as the proportion of their genomes shared through long IBD segments. These recent relatedness patterns differ markedly between African- and European-Americans: African-Americans show a distinct enrichment in South-to-North relatedness along the main historical migration routes.

**Figure 3:**
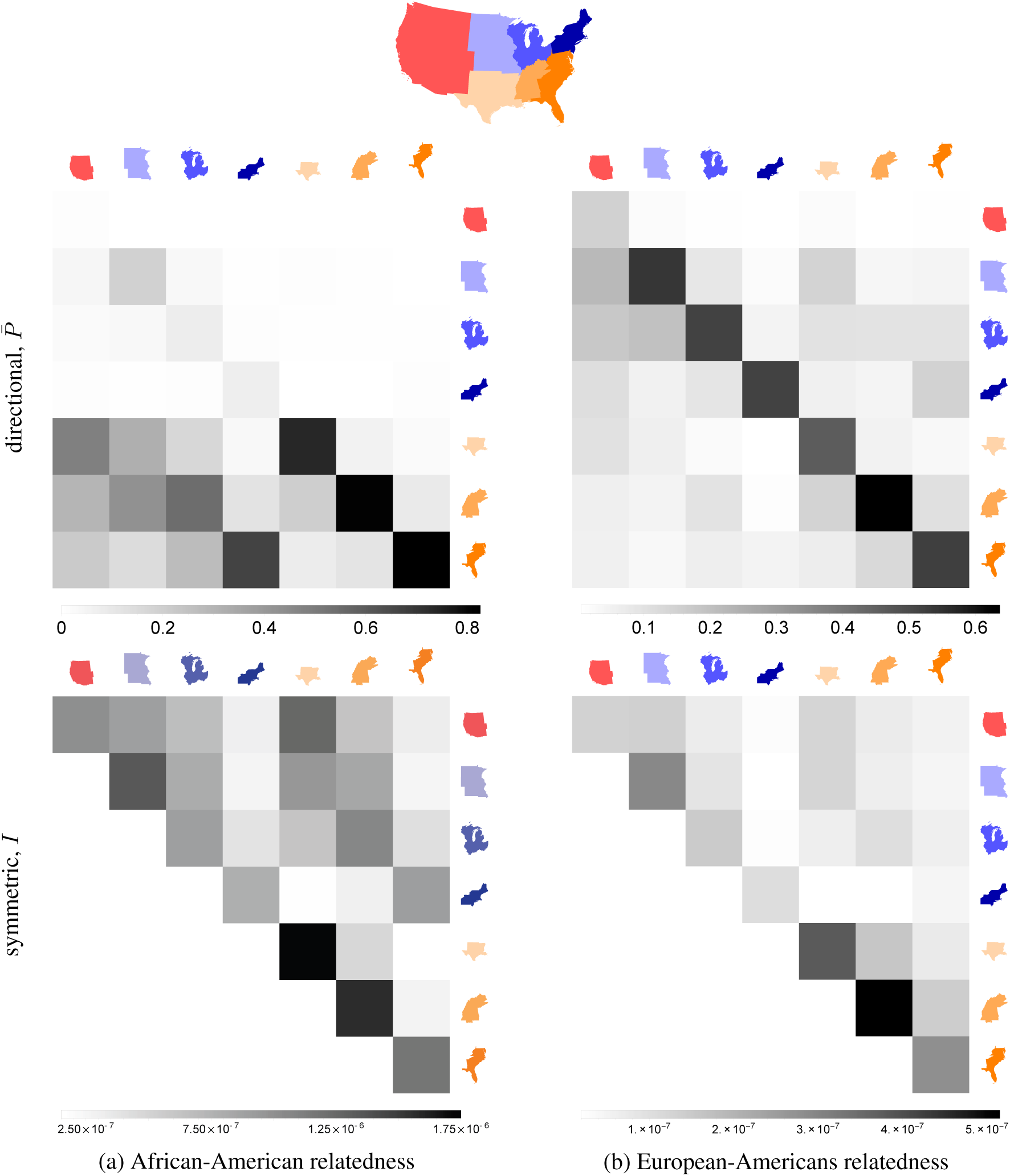
Pairwise genetic relatedness across US census regions among (A) African-Americans, (B) European-Americans, and (C) African-Americans and European-Americans. (D) Census-based prediction for African-Americans (see Methods). On each map, the thickness and opacity of a line connecting any two geographic regions show the relative strength of relatedness between those regions. Relatedness between regions with fewer than 10,000 possible pairs of individuals is not shown (see Methods for details). All numbers are in units of cM. (E) Decay of average IBD (shown in logarithmic scale) as a function of distance using IBD segments of length 18 cM or larger from HRS (dots), compared to the analytical model (lines).

To compare these relatedness patterns with recent migration data, we used the 20th century US census data and a simple coalescent model to estimate the expected relatedness between geographic regions (see Methods). Census-based predictions (Fig. 3D) are correlated with IBD-based observations (Fig. 3A) if we consider non-identical pairs of regions (Mantel test *p* = 0.019). Limiting the comparison to the South-to-North and South-to-West relatedness, to capture migration routes specific to the Great Migration, yields *p* = 0.063 (using the 2010 region of residence) and *p* = 0.015 (using place of birth) (see Methods).

Figures 3C and S24 show the relatedness between African-Americans and European-Americans. African-Americans across the US are more related to European-Americans from the South than to those from the North or West. In addition, European-Americans from the South tend to be more related to African-Americans in the North than to those in the South. This increased relatedness with increased distance is unusual in population genetics, but is easily explained: The ancestry-biased migration is also a relatedness-biased migration. The reduced relatedness between northern European-Americans and African-Americans may also be reinforced by recent European migration, because the new migrants were more likely to settle in the North but were less likely to be related to African-Americans.

## Fine-scale isolation by distance

Despite the unusual long-range relatedness patterns, identity-by-descent decays with distance within African-American communities in the South, reflecting isolation-by-distance (Fig. S5). To understand how migrations affect isolation-by-distance and identity-by-descent, we introduce a novel, simple model taking into account a diploid population density *n* and spatial diffusion constant *D*. In short, the displacement between parental birthplace and offspring birthplace of individuals is modeled as an isotropic random walk; the distribution of the times *t* to the most recent common ancestor of two individuals separated by distance *R* is calculated under a coalescent model; and the amount of genetic material shared IBD given a common ancestor at time *t* is computed as in (*24*). In this model, the expected fraction of genome shared IBD, through segments of length in *ℓ* = [*l*_min_, *l*_max_] (in Morgans), between two randomly chosen individuals separated by *R* has the simple form

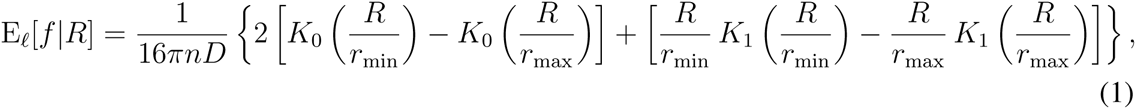

where 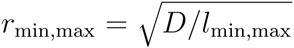, and *K_α_*(*x*) is the modified Bessel function of the second kind (see Methods).

Using IBD segments longer than 18 cM, we estimate population density *n*_AFR_ =1.9 km^−2^ and diffusion constant *D*_AFR_ = 63.5 km^2^/generation for African-Americans across the US, and *n*_EUR_ = 7.6 km^−2^ and *D*_EUR_ = 59.6 km^2^/generation for European-Americans (Fig. 3E). The ratio of European- to African-American population density is therefore 3.9. According to the 2010 US Census, 13% of the total population have self-identified as “Black or African American alone” and 72% self-identified as “White alone”. The ratio of European- to African-American population size from census is 5.5, in good agreement to our estimate above. Interestingly, the root mean squared displacement per generation, 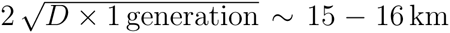, shows comparable local migration rates in European-Americans and African-Americans despite the different histories and population densities.

## Conclusion

The history of African-American populations combines strong ties to place with large-scale migrations (*4*). This comprehensive study shows the combined effects of fine-scale population structure, large-scale migrations, and admixture among African-Americans, giving us a better understanding of how the dramatic history of African-Americans shaped genomic diversity and, in particular, the sharing of haplotypes that can harbor rare, recent variation. From a medical genetics perspective, the sharing of recent haplotypes is a good proxy for the sharing of large-effect deleterious alleles (*25*). Rare genetic traits are therefore much more likely to be shared over long distances among African-Americans than among European-Americans, particularly along the routes of the Great Migration, but their spatial distributions over short ranges remain far from uniform. In addition, the ancestry-biased migration indicates a strong correlation between genetic and environmental population structure. Detailed demographic modeling should therefore inform the sampling and analysis of large genetic cohorts that include African-Americans.

## Acknowledgements

This work was supported by CIHR through the Canada Research Chair program and operating grant MOP-136855 (to S.G.) and NSF DMS award 1201234 (to C.D.B.).

## Supplementary Materials

### 1 Methods

#### 1.1 Data

We used the genotype data of 12,454 individuals from the Health and Retirement Study (26) (HRS), genotyped on the Illumina Human Omni 2.5M platform, and of 2,169 African-American individuals from the Southern Community Cohort Study (*27*) (SCCS), genotyped on either Illumina Human Omni 2.5M or Human 1M-Duo platforms. The HRS cohort includes 1,649 individuals who self-identified as African-Americans (non-ambiguously in both HRS Tracker and dbGaP databases) and 10,432 individuals who self-identified as European-Americans. There are also 366 individuals labeled as “Others” whom we have not used in our main analyses (except in a PCA analysis, discussed below). The remaining 7 individuals have ambiguous, non-matching race identifiers in HRS Tracker and dbGaP, and we have, thus, excluded them from our analyses.

We performed comparisons with data from 23andMe (12) and from 97 individuals of African ancestry from the southwest USA (ASW) from the 1000 Genomes Project^1^ (*28*). The 23andMe cohort includes many African-American individuals and has been the subject of a detailed population genetic analysis (*12*), and the ASW cohort has been a reference African-American population in recent studies. However, these two cohorts were not meant to be representative of the US population. The 23andMe database has a complex ascertainment scheme, which may cause biases in ancestry and socioeconomic status. In particular, biases in regional representation and a small amount of survey response errors might lead to a lower European ancestry proportion. These possible biases are described in detail in (*12*). Similarly, the ASW cohort was assembled from duos and trios with at least one Oklahoma resident, but with no attempt to reach geographic or demographic representativeness. For comparisons with the 23andMe study, we used the global ancestry proportions reported in (*12*), because the genotype data is not publicly available.

The use of these samples for the present study was approved by the IRB at McGill University and Stanford University, where the analyses were performed.

#### 1.2 Data merging and quality control

The HRS genotype data that we received had been already quality controlled, filtered, and phased. The SCCS cohort comprises data from 648 individuals in a breast cancer study (genotyped on Illumina Omni 2.5M platform) and 760 individuals in a prostate cancer study, 484 individuals in a lung cancer study, and 277 individuals in a colorectal cancer study (genotyped on Illumina Human 1M-Duo). All genotyped individuals were either cases or controls in their respective nested case-control studies. We converted the lung cancer dataset from human genome assembly hg18 to hg19 using the LiftOver utility from the UCSC Genome Bioinformatics Group and merged the four separate SCCS datasets into one using PLINK 1.9 (*29*). During the merge process, we removed markers to which more than one name was assigned at the same position along a chromosome; removed markers with missing genotype calls; corrected unambiguous strand misassignments and removed ambiguous strand (mis)assignments; removed multi-allelic markers; and, finally, filtered the data for missing calls (*30*) first based on genotypes (PLINK argument `--geno 0.0125`) and then based on call rates per individual and minor allele frequency (PLINK arguments `--mind 0.0125 --maf 0.01`). The final SCCS dataset contains 2,128 individuals and 585,527 variants after these steps. We then used the same process to merge the HRS data with those of SCCS and ASW, resulting in a single dataset in PLINK format with 14,679 individuals and 553,795 variants. Performing a PCA on the data (pruning for LD leaves 77,902 markers), we found no batch effects (see Fig. S13). We then phased the merged data with SHAPEIT2 (*31*) (default arguments), and converted the output to PLINK format (while preserving the phasing information) using genetic map information from the 1000 Genomes Project data^2^.

#### 1.3 Geographic information

Geographic information in HRS is usually provided in the form of US census regions and divisions. We have used these locales in the ancestry analyses. ZIP code information for HRS study participants is available, but use of this data is restricted. We used zip code data only for the fine-scale spatial analysis of identity-by-descent relatedness. For SCCS, latitude and longitude coordinates of clinics were available. In the IBD analysis, we assigned the ASW individuals to the West South Central census division. In terms of geographic locations, we restrict our analyses to the census divisions in the contiguous United States (i.e., Pacific, Mountain, West North Central, East North Central, Middle Atlantic, New England, West South Central, East South Central, South Atlantic).

For the individuals in HRS, we only consider the ones born in the contiguous US who, at the time of sampling in 2010, also lived in the contiguous US; this reduces our sample size in HRS to 10,974 individuals of which 1,501 are self-identified African-Americans and 9,308 are self-identified European-Americans (with the remaining individuals being classified as “Others”). There are 4 additional individuals satisfying the geographic constraints above but have discordant race identifiers in two different data files provided with the cohort data; these were removed from any downstream analysis. Among the unambiguous self-identified African-Americans and European-Americans mentioned above, there are respectively 10 and 427 individuals also self-identifying as Hispanics. The former 10 individuals are only included in our analysis of Hispanics status.

African-American sample sizes in the New England and Mountain census divisions are small. We therefore merged the New England and Middle Atlantic divisions, and considered the Northeast census region as a whole. Similarly, we merged the Mountain and Pacific, and considered the West census division as a whole. The total number of geographic locales under consideration was therefore 7: Northeast, Midwest consisting of 2 divisions, South consisting of 3 divisions, and West (see Table S5).

#### 1.4 IBD inference

We used GERMLINE (32) (arguments `-err_hom 1 -haploid -bits 32 -w_extend`) to infer IBD tracts of length 3 cM or larger shared between individuals from the HRS, SCCS, and ASW cohorts. GERMLINE is prone to false positive IBD assignment (see, e.g., (*33*)). We used a heuristic filtering scheme to identify and filter regions with large excess of IBD. We first count the number of overlapping IBD segments at each genomic position across all individuals. A chromosomal region is then marked as “forbidden” if the total number of IBD segments overlapping it is larger than 25,000, since the background IBD has an approximate depth of 15,000. Next, two forbidden regions will be merged as one if they are less than 0.1 cM apart. IBD segments that overlap these forbidden regions are excluded from the downstream analysis unless they extend outside the forbidden regions by at least 3 cM. In that case, we presume that there is sufficient evidence in the non-forbidden regions, and the segments are therefore kept. After this filtering process, we are left with 8,664,251 IBD segments out of the total of 71,633,425.

#### 1.5 Regional relatedness using genomic data

Geographic information along with the inferred IBD segments were used to construct a relatedness metric between individuals and geographic regions within our cohorts. We first bin the IBD segments by length. The first bin contains segments of length between 3 cM to 10 cM, the second bin contains segments from 10 cM to 18 cM, and the last bin contains segments of length 18 cM or longer. The latter bin corresponds to common ancestors living about 8 generations ago and is the focus of most of our discussion. Sorting the individuals by region and by African-American status within each region, we form two sparse relatedness matrices: 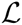 which contains the total IBD *length* shared between each pair of individuals, and 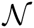 which contains the total *number* of shared IBD segments between each pair of individuals. We have to emphasize that the diagonal elements of 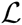 and 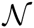, which represent self-IBD, are zero by definition.

We next remove the contributions of closely related individuals from these matrices as follows. The HRS study has already identified 89 pairs of individuals having kinship coefficients greater than or equal to 0.1. To be consistent with the definition from HRS, we used PLINK to calculate kinship coefficients for SCCS and ASW individuals, labelling individuals with kinship coefficient of 0.1 or higher as related individuals. We find 22 related pairs among SCCS individuals, 62 related pairs among ASW individuals, and 1 related pair between HRS and SCCS individuals (details below).

To see how geographic regions are associated based on the genetic relatedness of their inhabitants, we consider average pairwise IBD relatedness between regions (*23*). We take as the average pairwise relatedness *L* between two regions *R*_1_ and *R*_2_ the mean length of IBD segments shared between pairs of individuals, where one individual is from *R*_1_ and the other from *R*_2_. In addition, we consider the relationships between individuals of specific ancestry *S*_1_ and *S*_2_, each representing either African-American or European-American. Thus, the average total shared IBD length becomes

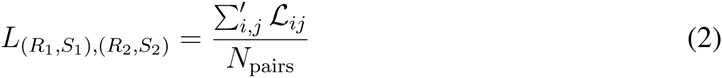

where

- *i* and *j* represent individuals as indexed in 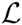;
- the primed sum runs over relevant pairs (*i*, *j*) such that *i* < *j*, (*R*(*i*), *S*(*i*)) = (*R*_1_, *S*_1_) and (*R*(*j*), *S*(*j*)) = (*R*_2_, *S*_2_), where *R*(*i*) and *S*(*i*) denote the region and race status for individual *i*;
- *N*_pairs_ = *n*_1_*n*_2_ if *R*_1_ ≠ *R*_2_ and *n*_1_(*n*_1_ − 1)/2 otherwise with *n_i_* being the number of individuals with attributes (*R_i_*, *S_i_*).

Using the metric defined above, we can calculate the pattern of relatedness between geographic locations with African-Americans, with European-Americans, and between African-Americans and European-Americans. The first two matrices are symmetric if we change the order of regions, whereas the last one is not.

##### 1.5.1 Visualization of regional relatedness

The following criteria was used for visualization of the IBD relatedness between regions. Due to the small number of African-American sampled individuals in the northern and western regions, the total number of IBD segments shared between these regions is small compared with that between other regions: see the bottom row in Fig. S22. Relatedness estimations are noisy for such pairs, and a scale that accommodates these noisy results would not allow for detailed comparison of less noisy results. Therefore, in Figs. 3, S20, and S21, we did not draw the lines between any two regions for which the total number of possible pairs of IBD individuals is less than 10,000 (e.g., notice the lack of connecting lines from West North Central to West). Since a significant number of the individuals in HRS are European-Americans, the number of IBD segments shared between European-Americans residing in any two regions is large enough to ensure the significance of the results, even when we restrict the analysis to the longest IBD segments (see the bottom row in Fig. S23).

#### 1.6 Regional relatedness using census data

The relatedness pattern derived from genomic data can be compared with historical migration records, available from Integrated Public Use Microdata Series (IPUMS) (2). We downloaded census data from 1900 to 1980 and extracted census year, census region, age, race, birth place, and weighted representation of each sample; the latter is the number of people in the population represented by the sampled individual. We focus on the people in the age group of 20- to 30-year olds for any decade, and consider the migrations of African-Americans and European-Americans separately. We assume a generation time of 30 years, thereby taking census years 1900, 1910, and 1920 as generation 3; 1930, 1940, and 1950 as generation 2; and 1960, 1970, and 1980 as generation 1. For each race group, we construct a matrix whose elements *m_ij_* are the number of migrations for each generation from region *i* to region *j*; this matrix is highly asymmetric because of asymmetric migrations.

We now construct a heuristic census-based measure of relatedness between regions. Let us define 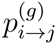 as the proportion of individuals at generations *g* − 1 in region *j* with ancestors at generations *g* in region *i*. In other words, the (*i*, *j*) element of the matrix *P*^(^*^g^*^)^ is

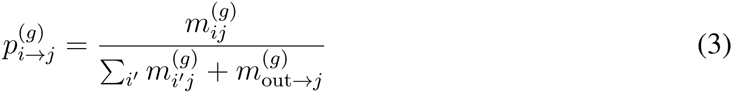

where *g* ∈ {1, 2, 3} denotes the generation time of the ancestral population, and 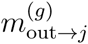 is the number of migrants from outside of contiguous United States into the census region *j*. Had we not included migrations from outside US into the mainland US, *P*^(^*^g^*^)^ would have been column-normalized (i.e., normalized with respect to the destination census regions).

We construct a three-generation transition matrix as:

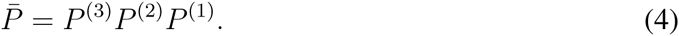

This definition for 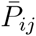 takes into account all possible migration routes starting at region *i* and ending at region *j* that could have taken place in the span of the three generations.

To estimate genetic relatedness between different geographic regions, we further make the extremely coarse assumption that population sizes were constant before 1910, and that populations were randomly mating. These assumptions allow us to model expected relatedness within regions using coalescent theory before the massive 20th century migrations. Neither assumption is expected to hold, but the resulting relatedness metric remains informative as long as census population size correlate with the expected time to the most recent common ancestor for a given pair of lineages in a region.

Given 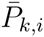 as the probability of a sampled individual from region *i* having an ancestor from region *k*, we define the census relatedness metric between regions *i* and *j* as

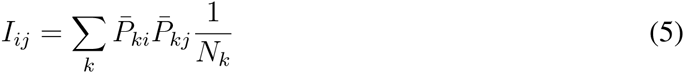

where *N_k_* is the census population size of region *k*. Population size matters, because in larger populations, it is less likely that a given pair of individuals share a common ancestor. If *N_k_* is large enough, the number of common ancestors at each generation is inversely proportional to *N_k_*, and therefore the expected recent shared ancestry is approximately inversely proportional to *N_k_*. Thus, *I_ij_* is proportional to the probability of two individuals from regions *i* and *j* having ancestors from (any) region *k* times the probability that these ancestors have a recent common ancestor within region *k*. Unlike 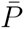 which is a directional metric, *I* is non-directional and symmetric and can be directly compared with genetic relatedness matrix *L* in Eq. (2), which was estimated using IBD data. The regional relatedness patterns derived using 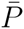 and *I* are shown in Fig. S4.

**Figure 4:**
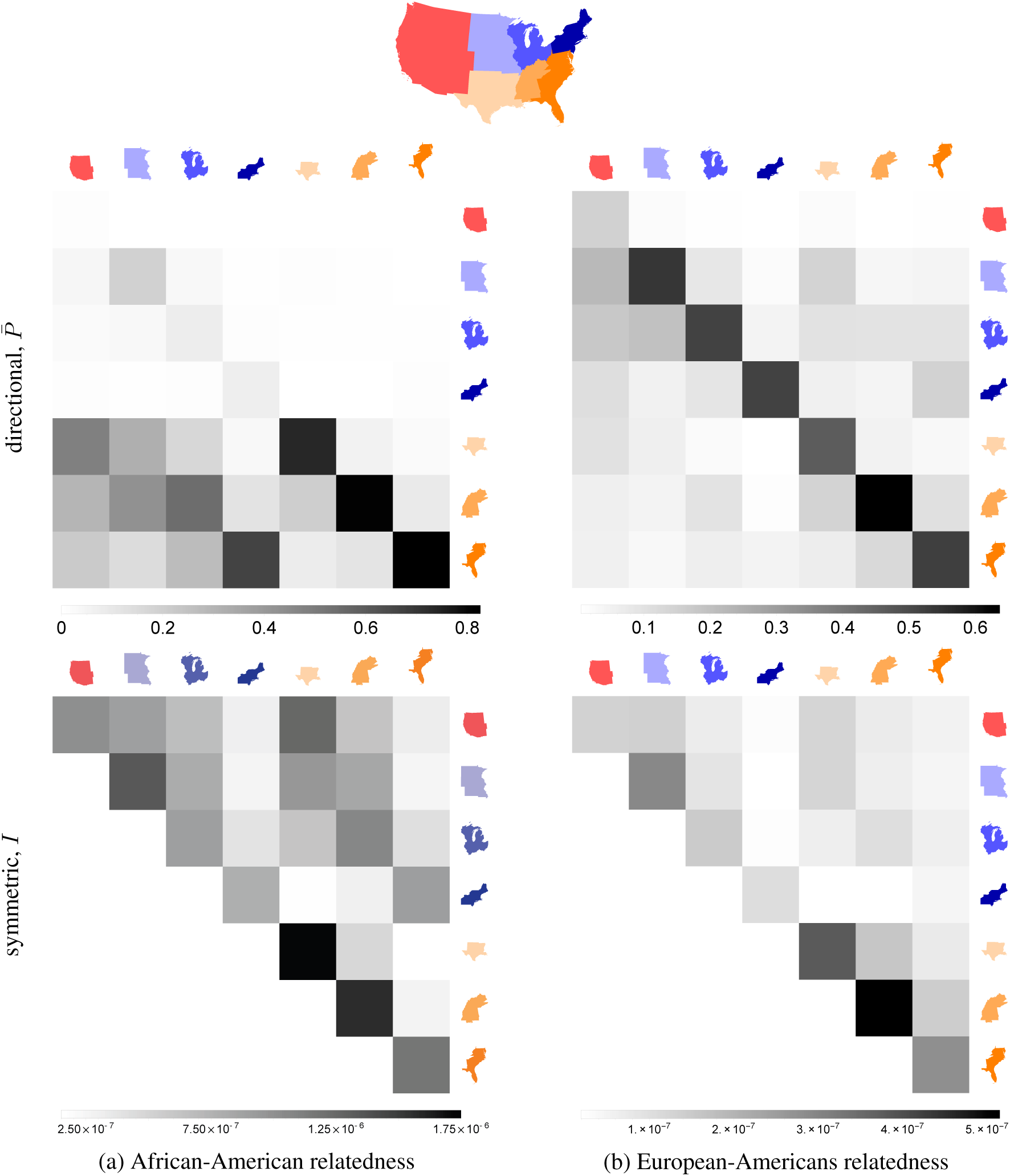
Census-based predicted relatedness between (a) African-Americans and (b) European-Americans across the US census regions. The top row shows the directional metric 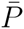, whereas the bottom row shows the symmetric one *I*. In the top figures (read column-wise), each column shows for its respective census region the proportion of ancestral population which originated from other census regions. See Fig. S25 for the numerical values of these regional relatedness metrics.

#### 1.7 Significance test for genomic versus census-based relatedness

To test the hypothesis regarding South-to-North migration corridors, we consider the matrix elements corresponding to relatedness between the three southern regions (South Atlantic, East South Central, West South Central) and the three northern ones (Northeast, East North Central, West North Central), forming a 3 × 3 matrix from the census data to be compared with the corresponding matrix from IBD data. To quantify the correlation between these two matrices, we use the Mantel test as follows. Given that all the elements in each of these two matrices are independent, we perform 9! possible permutations on the elements of the matrix derived from the census data and calculate the Pearson correlation coefficient between the original IBD matrix and the permuted census matrix. We then accept or reject each permutation based on whether the calculated correlation coefficient is lower or higher than the correlation coefficient between the two original (non-permuted) matrices. The *p*-value is given by the ratio of the number of rejections to the total number of permutations (see main text for the numerical values). The *p*-value reported in the main text for the relatedness between South to North and West are derived by performing a random subset of 10^7^ permutations out of a total of 12! ones.

**Table 2:**
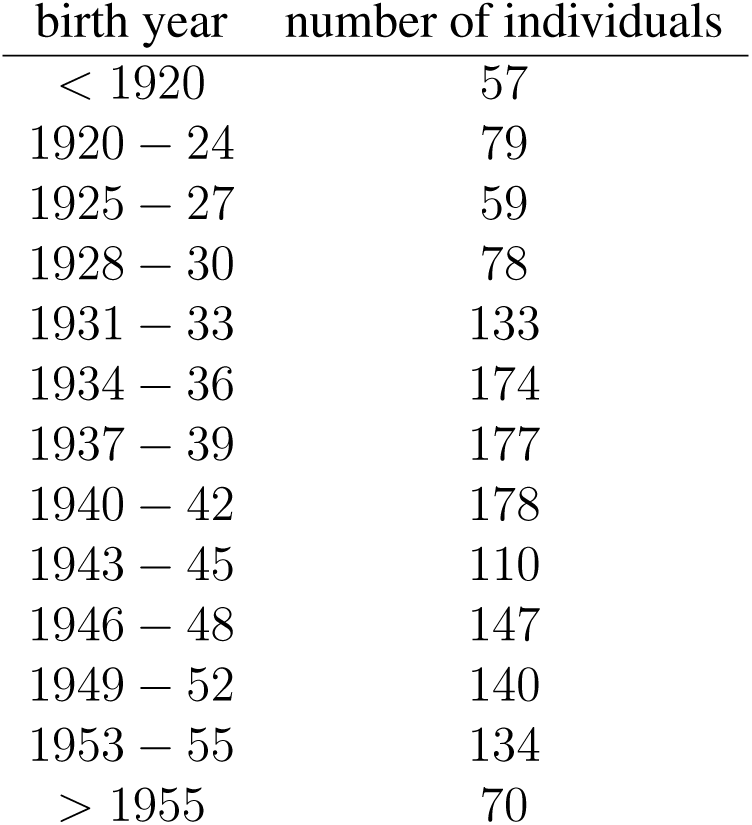
Distribution of birth years in HRS

We also perform this test while using the region of birth of HRS individuals as their location. Given the average year of birth (1939.8) and the birth year distribution (Table S2) in HRS, we only take for consistency generation 3 from the census data (see definition above) and write 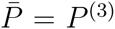 as our overall directional relatedness matrix (compare with Eq. (4) above). We then proceed as before to calculate the non-directional (symmetric) relatedness *I*. Given the new census-based prediction (using only *g* = 3 above) and the IBD relatedness pattern (using the region of birth), we perform a Mantel test as before in order to find the correlation between the data and our prediction.

#### 1.8 Relatedness and isolation-by-distance

We wish to model the expected IBD relatedness between individuals in a spatially extended population. Our starting point is an idealized population living on a set of islands (or demes), with random mating within islands and migrations between the islands. We will consider a limiting example of a continuous population below.

We are first interested in the probability that a genomic segment of given length, stretching across a specific locus, is shared identical-by-descent between two randomly selected individuals living on different islands. For identity-by-descent to occur, we need two events to happen: first, lineages at that locus must have coexisted on one unknown island at some point in the past. Second, these two geographically coexisting lineages must also have coalesced further in the past.

We measure time in generations and track lineages backwards in time. At each generation, we assume that the displacement between parental birthplace and offspring birthplace follows a random walk. Each lineage traverses follows a random walk on the islands, with each step representing one generation back in time, connecting an individual to the ancestor from whom the locus is inherited. The lineages are then traced back until the time at which both ancestors coexist on the same island and coalesce in the most recent common ancestor in the next step back in time. We can, therefore, symbolically write for the total probability of coalescence at a given generation as a probability of coexistence times a probability of coalescence:

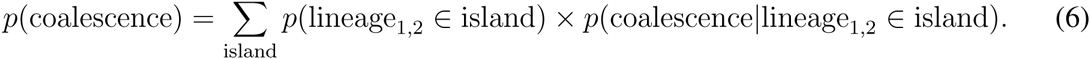

To derive the probability of coexistence, we first want to estimate the expected position of a lineage given its position in the past. Concretely, let x_0_ be the current location of an individual at *t* = 0. We would like to find Φ(x, *t*|x_0_), the probability that an individual’s lineage is on island x at *t* generations ago, given that it is currently on island x_0_.

By construction, the probability Φ(x, *t*|x_0_) takes into account contributions from *all* possible space-time paths that start at x_0_ and end at x at time *t*. For instance, a possible path is to arrive at x at *t*/2 and stay at that position until *t*, whereas another path is to arrive at x at *t*/3, leave x at the next step for a series of random walks to finally arrive at x again at *t*.

Consider a region of area Δ*A_i_* that encompasses a deme with haploid population 2*n*(x*_i_*, *t*)Δ*A_i_*, where *n*(x*_i_*, *t*) is the effective diploid population density at position x*_i_* and time *t* in the past. The probability of that two lineages in Δ*A_i_* coalesce in a given generation is

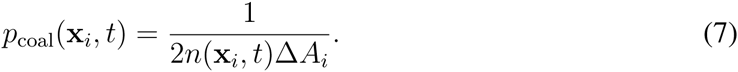

This expression does not consider the possibility of multiple coalescent events and is therefore appropriate only for a number of generations that is much less than the population size. Moreover, the discrete probability of two lineages having coexisted on the deme at x*_i_* at time *t* in the past, given that they are a distance R apart (at x_0_ and x_0_ + R) at present (at *t* = 0), is

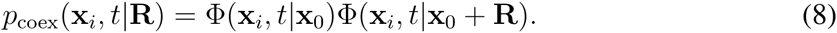

Therefore, the total probability of having a common ancestor *t* generations ago in the discrete model is

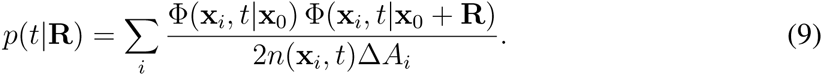

To go from a discrete random walk to the continuous limit, we set Φ(x, *t*|x_0_) → *φ*(x, *t*|x_0_)Δ*A*, where *φ*(x, *t*) is now a continuous probability density. Thus, in this limit, i.e. with ∑*_i_* Δ*A_i_*& → *∫* d^2^x &, we get

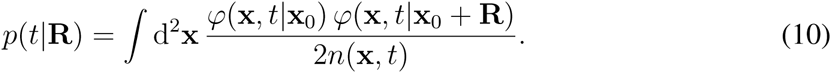

The continuous limit of a random walk process is the diffusion model. In this model, the probability density *φ*(x, *t*) of finding a lineage at an infinitesimal area d^2^x centered around x at generation *t* in the past obeys the two-dimensional partial differential equation

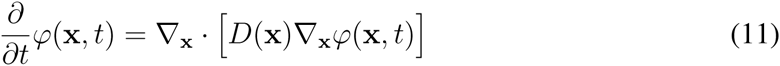

where the diffusion coefficient *D*(x) encompasses the information related, in the discrete model, to probabilities of taking a step to an adjacent island or staying on the same island^3^. Solving for *φ*(x, *t*|x_0_) amounts to solving equation (11) with initial condition *φ*(x, *t* = 0) = *δ*(x − x_0_) where *δ*(x) is the (two-dimensional) Dirac delta function.

For simplicity, we consider random walks with uniform probability of transitioning to any nearest-neighbor island, which translates to a constant (position-independent) *D* in the continuous model. We also assume that all islands have the same constant population size, leading to a population density which, on average, is constant in the continuous model. At each time step, the following processes occur for each individual: (a) reproduction and subsequent death according to the Wright-Fisher model and (b) migration of all, some, or none of the individual’s offsprings to other islands. With this definition, tracing back a lineage includes, in each time step, an offspring-to-parent generation and, potentially, a coalescence event with another lineage.

Under these assumptions, we have

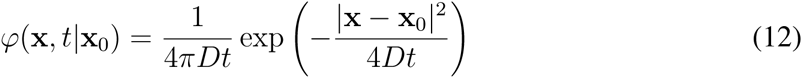

which, in turn, leads to

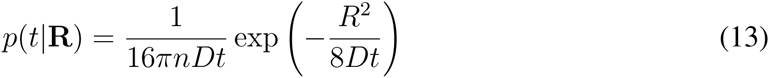

with *R* = |R|.

Next, following Palamara *et al.* (*24*), we have for the expected fraction of the genome shared through segments in the length range *ℓ* = [*l*_min_, *l*_max_] (in units of Morgans)

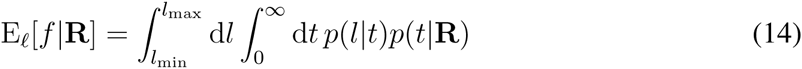

with *p*(*l*|*t*) = (2*t*)^2^*l* exp(−2*tl*) the probability density of an IBD segment of length *l* (in units of Morgans) spanning the locus shared by the two randomly chosen individuals whose lineages coalesce *t* generations ago. Performing the integrals above leads to the following closed form solution for the expected fraction of the genome shared as a function of spatial separation

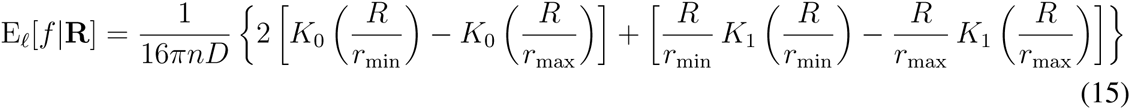

where *K_α_*(*x*) is the modified Bessel function of the second kind (*34*), and 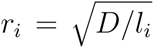 with *i* ∈ {min, max}. Expanding for small *R*, we find 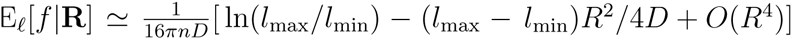 for small *R*.

We can use this estimator to find the shared fraction per chromosome; that is, for each chromosome, we set *l*_max_ in Eq. (15) equal to *L_c_* (the length of chromosome *c*) to get 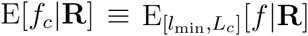. The total length of shared IBD tracts (across all chromosomes) between a random pair of individuals, therefore, becomes 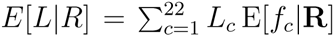; this quantity can be directly compared with that calculated from the IBD data to estimate the parameters of the model.

Access to the exact location of clinics at which the SCCS cohort was sampled allows us to investigate the relation between IBD relatedness and spatial distance. Having inferred possible IBD segments using GERMLINE, we calculate, for each pair of individuals from SCCS, the total length of shared IBD and the distance between the clinics in which they were sampled. We make the underlying assumption that each individual lives close to the clinic at which he or she was sampled. Each pair is then placed, based on the distance between the two individuals, into one of the length bins in {[0,1), [1,101), [101, 201), [201, 301),&} (all numbers in kilometers). The first length bin, [0, 1), contains individuals sampled at the same clinic. For each bin, we calculate the average pairwise IBD length (the sum of the IBD lengths of all pairs divided by the total number of points in the bin) and assign it to a distance equal to the midpoint of the bin (e.g., for the length bin [1, 101), the assigned distance is 51 km). The result is shown in Fig. S5.

Apart from the expected decay of relatedness with distance, we also notice the presence of a constant “background” IBD. This background IBD is larger for smaller IBD segments. As mentioned in the main text, this could be attributed to two possible factors: (a) GERMLINE has a higher false positive detection rate for shorter IBD segments (*33*) which is independent of the distance between individuals, or (b) smaller IBD segments, being much older on average, reflect history prior to migrations from Europe and Africa into the Americas. Since this relatedness patterns extends over long distances with little evidence for decay, we suppose that it is either due to false positives, or that there was enough mixing in the travels in to the Americas that present-day proximity is a relatively poor proxy for the proximity of ancestors prior to transatlantic travels. In either case, the background IBD can be modeled by adding a constant term to our model, representing the expected fraction of the genome shared IBD by individuals over long distances.

**Figure 5:**
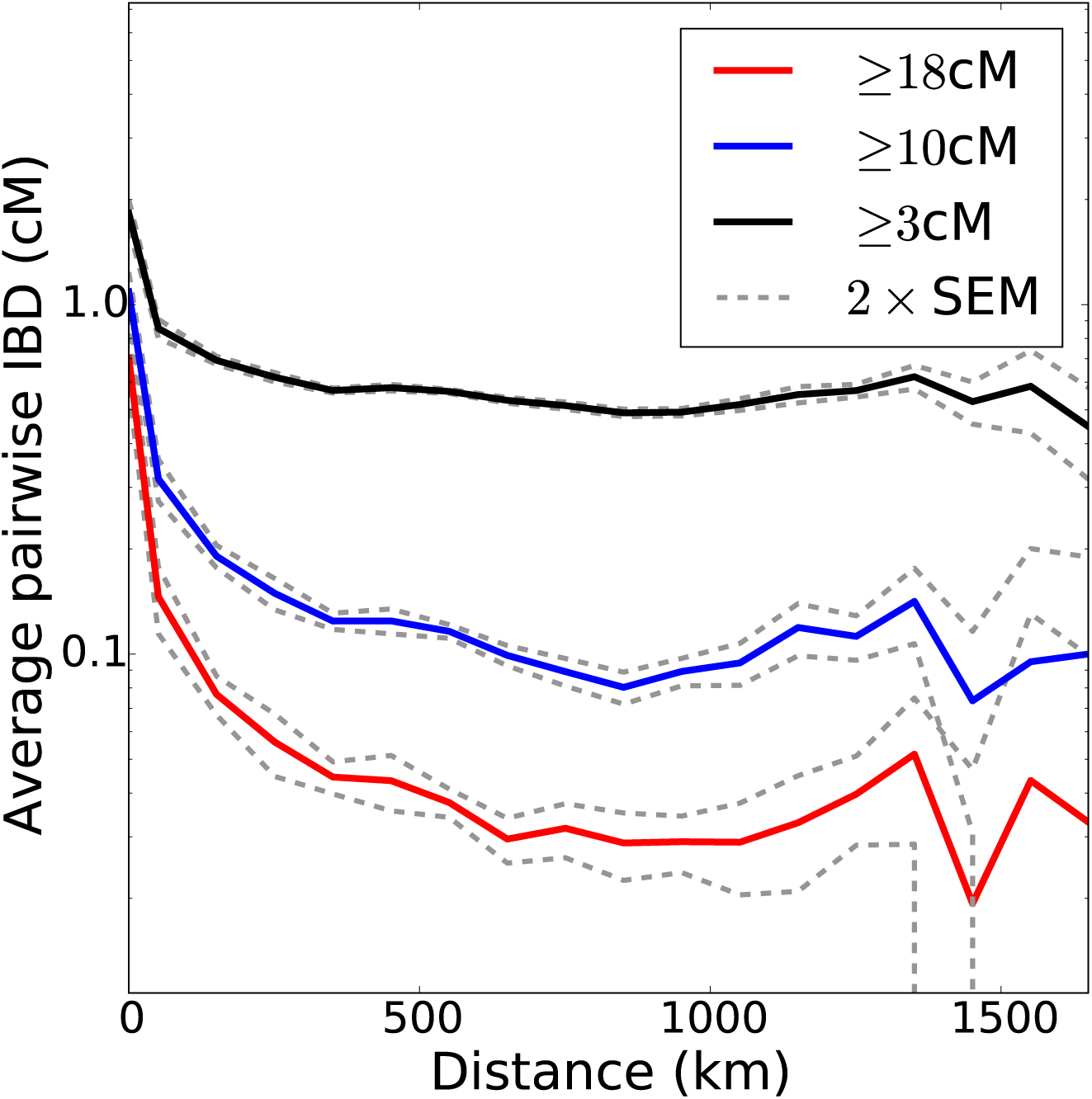
Decay of IBD sharing with distance, calculated for the SCCS cohort, for IBD segments of length 3 cM or larger (top), 10 cM or larger (middle), and 18 cM or larger (bottom). The plot is in log-linear scale, and the dashed lines represent two standard error deviations from the mean for the corresponding curve. The sharp fall-off of the dashed lines at large distances is due to the logarithmic scale of the vertical axis.

The parameters to be inferred in this model are the haploid population density *n*, the diffusion coefficient *D*, and background IBD *b*. By fitting the SCCS African-American IBD data for the 18 cM case (corresponding to the most recent sharing events), we find the estimated values *b*_18_ = 0.0389 cM, *n*_18_ = 2.8 km^−2^, and *D*_18_ = 88.6 km^2^/generation. The root mean squared displacement for African-Americans in the South is thus estimated, using the IBD data from SCCS, to be 18.8 km. We can use the population density and diffusion coefficient derived above to predict IBD decay for IBD segments of different lengths and estimate the background IBD for the other two cases (bins with segments of length 10 cM or larger and with segments of length 3 cM or larger), finding *b*_10|18_ = 0.120 cM and *b*_3|18_ = 0.546 cM. The resulting fits Fig. S6 show good agreement with the data.

#### 1.9 Expected *T*_MRCA_ given length of IBD segments

For reference, we derive the expected generation time to the most recent common ancestor (MRCA), given an IBD tract of certain length. The probability density of having an IBD segment of length *l* (in units of Morgans) spanning a chosen marker (denoted by *ζ*) inherited from a MRCA living *g* generations ago (assumed continuous for simplicity) is (*24*)

**Figure 6:**
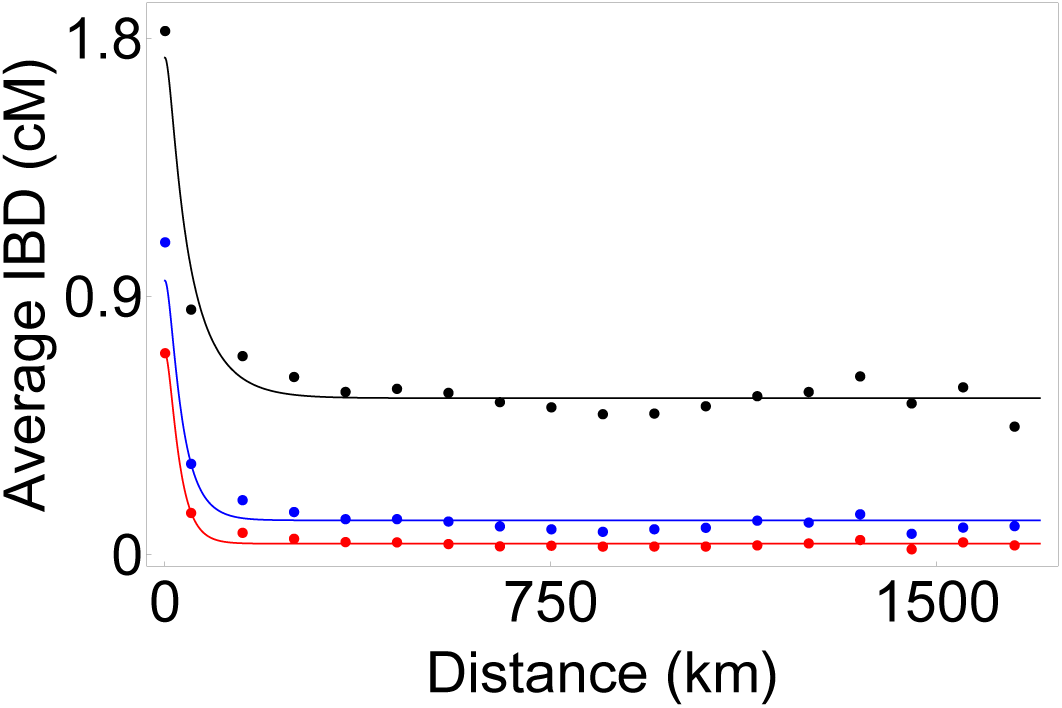
Estimated decay of IBD sharing with distance for IBD segments of length 3 cM or larger (top), 10 cM or larger (middle), and 18 cM or larger (bottom). Points represent the data and lines represent the model.

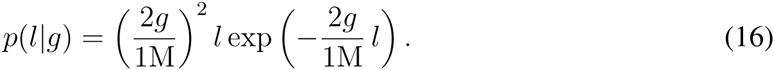

In the Wright-Fisher model, given the shared locus *ζ*, the probability of having a MCRA *g* generation ago is

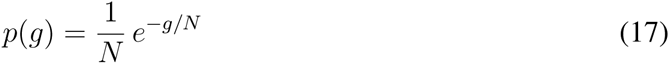

where *N* is the (constant) effective haploid population size. Therefore, given the length *l* of an IBD tract (in units of Morgans), we use (16) and (17) to find the expected value for the generation time of the MRCA

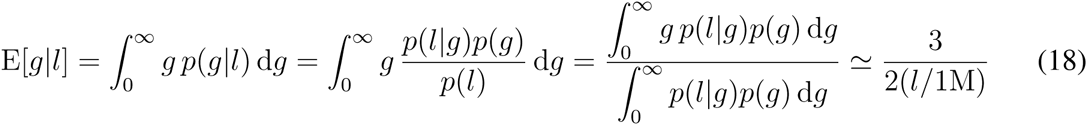

where we have assumed that the haploid population size *N* ≫ 1 in the last step.

#### 1.10 Local and global ancestry analysis

After the phasing process (discussed previously), we used RFMix (*11*) with arguments `PopPhased -- skip-check-input-format` for local ancestry inference along the genome. We used available parents among the trios in the Southern Han Chinese (CHS), Yoruba in Ibadan, Nigeria (YRI), and Utah Residents (CEPH) with Northern and Western European Ancestry (CEU) populations from the 1000 Genomes Project^4^ as a reference panel, comprising 50 CHS, 97 YRI, and 91 CEU individuals. We extracted the intersecting set of SNPs between our merged dataset and the three reference populations mentioned above, which we used as the input to RFMix. RFMix assigned continental ancestry of each marker in each sample to either CHS, YRI, and CEU, which we interpret as Native American/Asian, African, or European.

#### 1.11 Quality control of local ancestry inference

To ensure that the inferred low percentages truly reflect Native American ancestry, and not mis-assignment of European or African ancestry segments, we performed simulations based on a two- and a three-population admixture model. In both cases, we generated ancestry tracts for 50 admixed diploid genomes in a forward Wright-Fisher with a single pulse of admixture 8 generations ago.

For the two-population admixture model, the ancestry proportions in the simulated individuals were 74.96% African and 25.04% European. We copied genotypes from one YRI sample into European ancestry segments and one TSI sample into the African segments (both samples from the 1000 Genomes Project) to generate 100 haploid chromosome 1’s. Each chromosome 1 was generated using a distinct source chromosome in the YRI and TSI population. We then inferred the ancestries of the individual i (corresponding to haplotypes 2*i* and 2*i* + 1) with panels composed of samples chosen from 91 CEU, 50 CHS, and 96 YRI, ensuring that the individual from whom the genotypes were copied was not used in the reference panel. We inferred 74. 96% African, 24.95% European, and 0.09% Native American ancestry.

For the three-population admixture model, we simulated a sample of 100 haploid chromosomes with 80.9% YRI, 18.2% TSI, and 0.91% JPT ancestry, using the same method described above. In this case, the inferred proportions were 80.9% African, 18.2% European, and 0.94% Native American. These results are consistent with previous estimates of false assignment using a similar pipeline (*11*).

We also considered whether the amount of Native American ancestry in real samples correlated with the amount of European ancestry. If European segments are more likely to be misinterpreted as Native American, we would expect a positive correlation between inferred Native American and European proportions. Conversely, if the increased diversity in African segments led to higher rates of misidentification as Native American ancestry, we’d expect the correlation to be negative. The relation between Native American ancestry and European ancestry within SCCS is shown in Fig. S7. Within the southern states, only Louisiana shows a significant correlation. The lack of global correlation between European and Native American ancestry helps support the correctness of the inference.

Finally, we also compared global ancestry proportions inferred by RFMix and by ADMIXTURE (in supervised mode) and found an extremely high correlation between the estimates from the two methods, as shown in Fig. S8.

#### 1.12 Tracts and timing estimates

To infer time of admixture between ancestral populations and to identify migration models that give rise to the observed genome-wide patterns of ancestry, we use Tracts (*16*). We excluded for this analysis HRS African-Americans from non-mainland US (96 individuals), African-Americans with self-reported Hispanic ethnicity (32 additional individuals), and one additional African-American who was listed as “White, non-Hispanic” in HRS Tracker but as “African-American” in dbGaP. All individuals were kept in the other cohorts.

**Figure 7:**
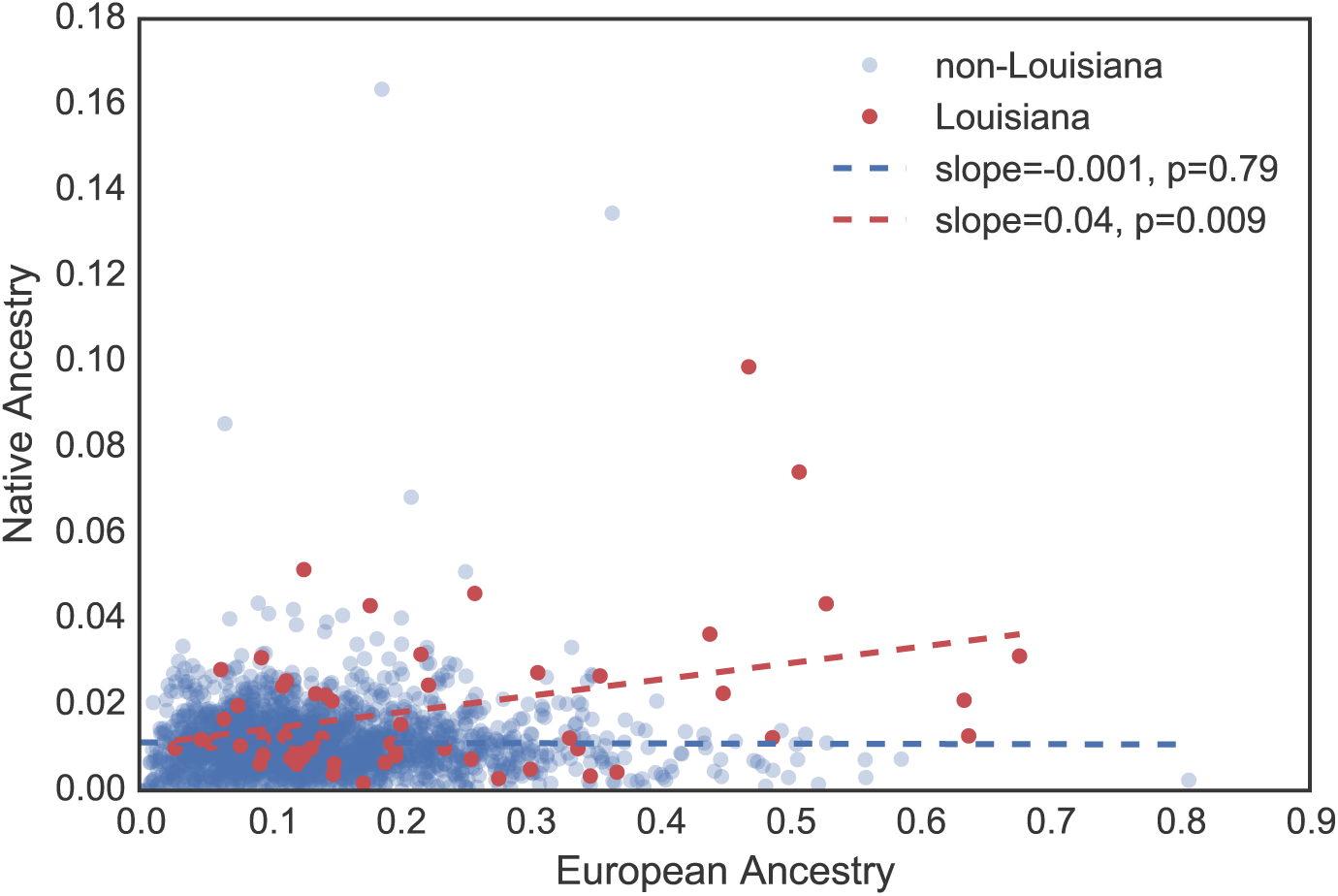
Inferred Native American versus European ancestry in the SCCS cohort.

**Figure 8:**
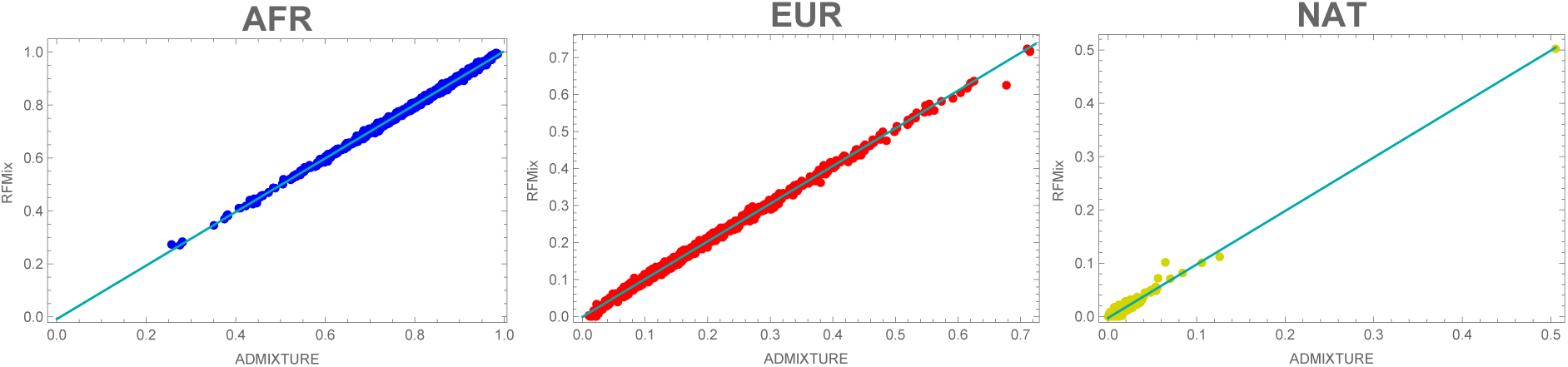
Correlation between continental ancestry (African, European, and Native American/Asian) estimates from RFMix and ADMIXTURE for HRS African-American individuals.

Optimization was performed for 6 models in each cohort: 2 two-population models, and 4 three-population models.

##### 1.12.1 Two-population admixture models

The first two-population model, `pp`, consists of a single pulse of discrete admixture between an African (AFR) and a non-African (NONAFR) population. The second model, `pp_xp`, considers a pulse of admixture from each population, followed by a second pulse of admixture from the NONAFR one. In the model nomenclature, migration events are described by strings separated by underscores. Each string has one letter per population, with `p` indicating a pulse of migration from the respective source population, and `x` indicating no migration from that population. For example, the model `pp_xp` has two events; the first event, `pp`, has discrete contributions from populations 1 and 2, and the second event, `xp`, has a contribution only from population 2.

Optimization for each model was performed using a brute force search over a grid of parameter points, followed by a local refinement from the maximum likelihood grid-point. Segments below 11.7 cM (corresponding to the first two bins in our histogram) were not used in the optimization process lest their numbers might have been less accurately estimated. However, model predictions for these segments were reasonably accurate for all models and cohorts. In the likelihood optimization, the total ancestral proportions for the population were held fixed; the optimization was performed over the timing of the admixture events and the relative contributions of the distinct pulses of admixture from the same source. The resulting histories and corresponding likelihoods are shown in Fig. S9.

**Figure 9:**
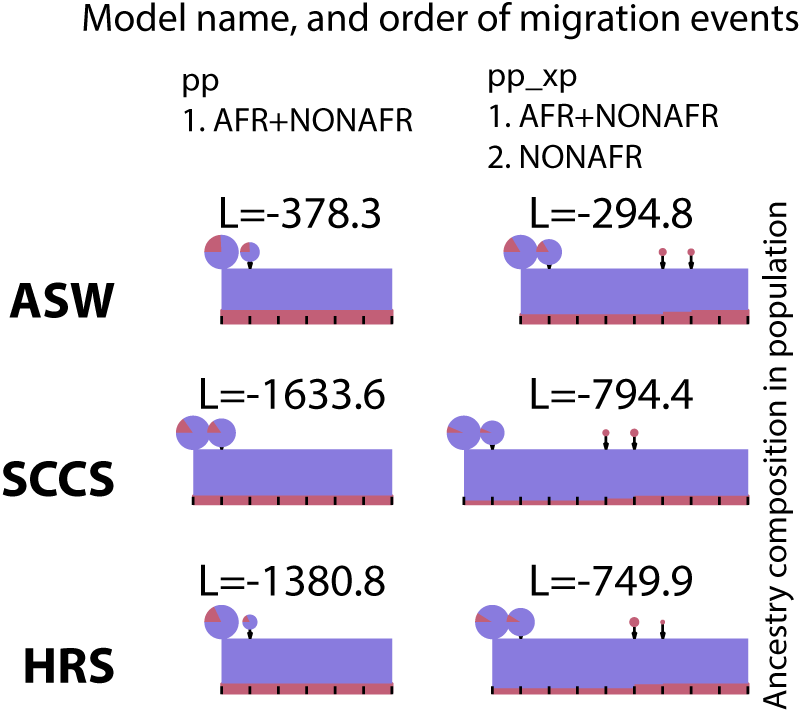
Estimated histories for two-population models, with the corresponding log-likelihoods. African ancestry is displayed in blue, and non-African ancestry in red. Rectangles show the proportion of each ancestry at each generation. Pie charts represent migrations, with the size of the pie representing the amounts of migrants at a given generation, and the sectors represent the proportion of migrants coming from each source population.

In addition to the global ancestry proportions, the `pp` model has a single free parameter (the timing of admixture), whereas the `pp_xp` model has three (two times of admixture and the relative contributions of the first and second non-African admixture). The `pp_xp` model outperforms the `pp` model by 631 log-likelihood units in the HRS, and by 839 log-likelihood units in the SCCS. We can reject the `pp` model according to either the Akaike information criterion or the Bayesian information criterion with *n* = 100 data points (one point per bin and per population).

##### 1.12.2 Three-population admixture models

In the three-population case, the `pxp_xpx` model consists of a founding admixture of African and Native American migrants, followed by a subsequent pulse of European admixture. The `ppp_xpx` model consists of a founding admixture event involving the three populations, followed by a subsequent pulse of European admixture. The `pxp_xpx_xpx` model has a founding admixture of African and Native American ancestors, followed by two pulses of European admixture. Finally, the `xpp_pxx_xpx` model has a founding population of Europeans and Native Americans, followed by a pulse of African admixture, followed by a pulse of European admixture. These histories are shown in Fig. S10. The best-fit model for the SCCS and HRS is the `pxp_xpx_xpx` (see Fig. S11 and S10).

**Figure 10:**
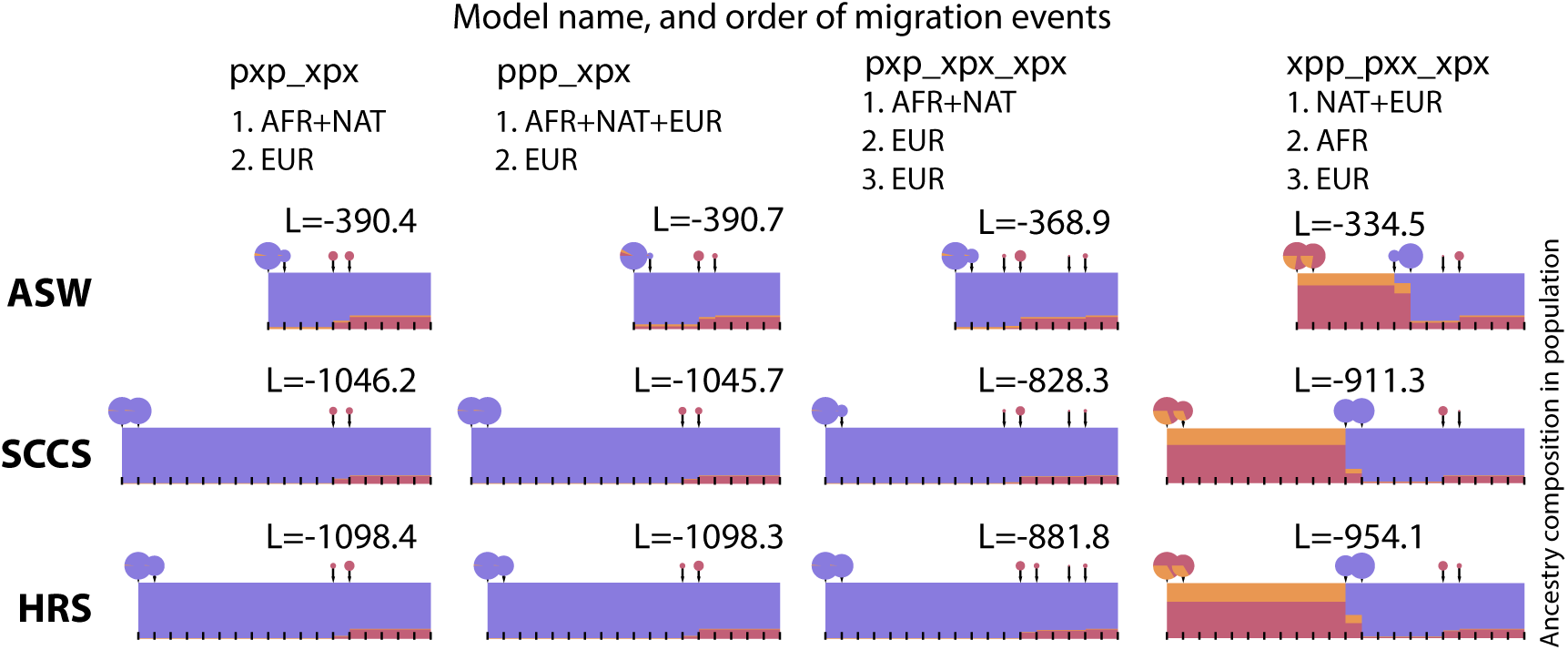
Estimated histories for three-population models, with the corresponding log-likelihoods. African ancestry is displayed in blue, and non-African ancestry in red. Rectangles show the proportion of each ancestry at each generation. Pie charts represent migrations, with the size of the pie representing the amounts of migrants at a given generation, and the sectors represent the proportion of migrants coming from each source population.

In addition to the three global ancestry proportions, model `pxp_xpx`, has two free time parameters. Model `ppp_xpx` has three parameters (two times of admixture and one relative contribution between the first and second pulse). Models `xpp_pxx_xpx` and `pxp_xpx_xpx` each have four parameters (three times and one relative contribution). For the HRS and SCCS datasets, the `pxp_xpx_xpx` model has the best likelihood. Since it outperforms the simpler models by 200 log-likelihood units, it is supported by either the Akaike information criterion or the Bayesian information criterion with *n* = 150 data points.

**Figure 11:**
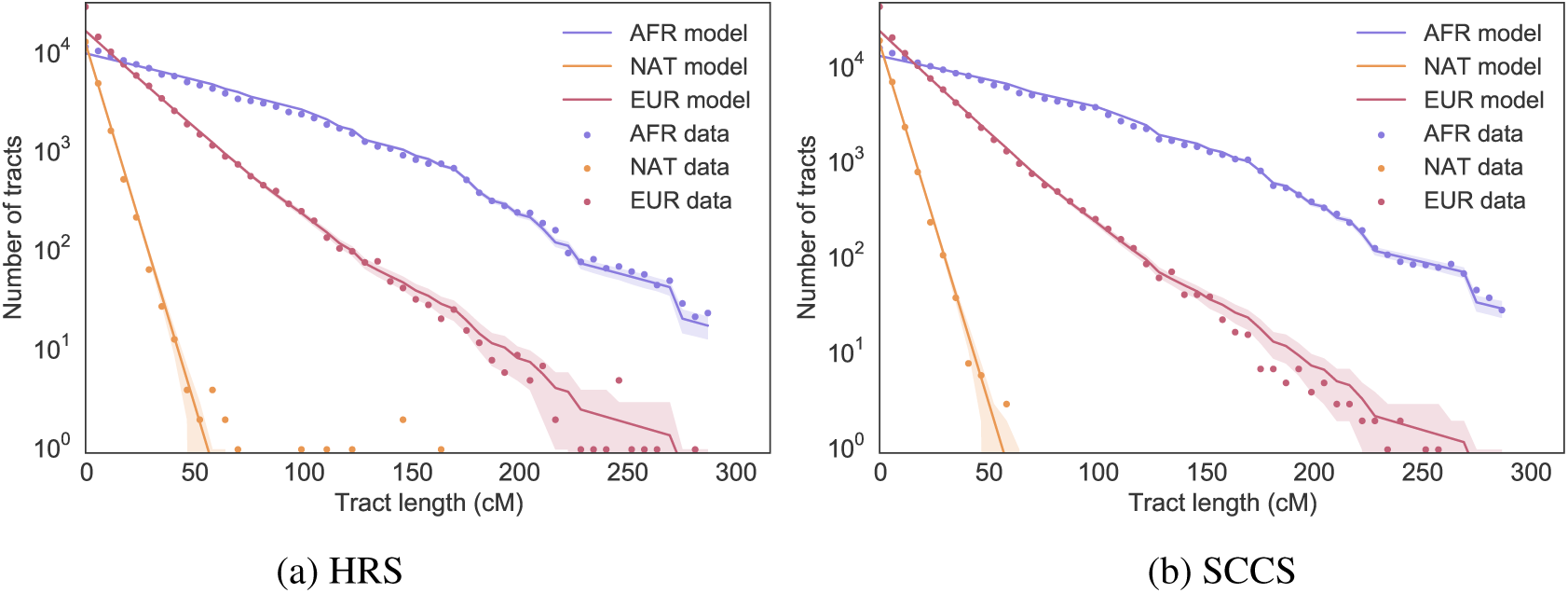
Comparison between observed tract length distribution (dots) and expectation under the best-fitting model (solid lines) for the HRS (a) and SCCS (b). Shaded areas represent one standard deviation departures from model expectations.

##### 1.12.3 Confidence intervals for timing estimates

Confidence intervals for all parameter values were obtained via bootstrap (Table 3). For each model, we generated 100 bootstrap populations by resampling individuals with replacement. We performed parameter inference for each bootstrap population, and computed the 95% confidence interval of the resulting distribution of parameters. These confidence intervals account for the finite number of individuals in the sample. However, they do not account for biases resulting from population structure or model mis-specification. Because of the large sample size, these biases are likely more important than the uncertainty measured by the bootstrap.

### 2 Supplementary text, figures, and tables

#### 2.1 Additional details on cohorts

#### 2.2 Number of individuals in each census region or division

Movement of HRS individuals between their time of birth and the 2010 sampling year is represented in Fig. S12. Because of these migrations, the number of individuals in each census region or division depends on whether we assign each individual to their region/division of birth or region/division of residence in 2010.

**Table 3:**
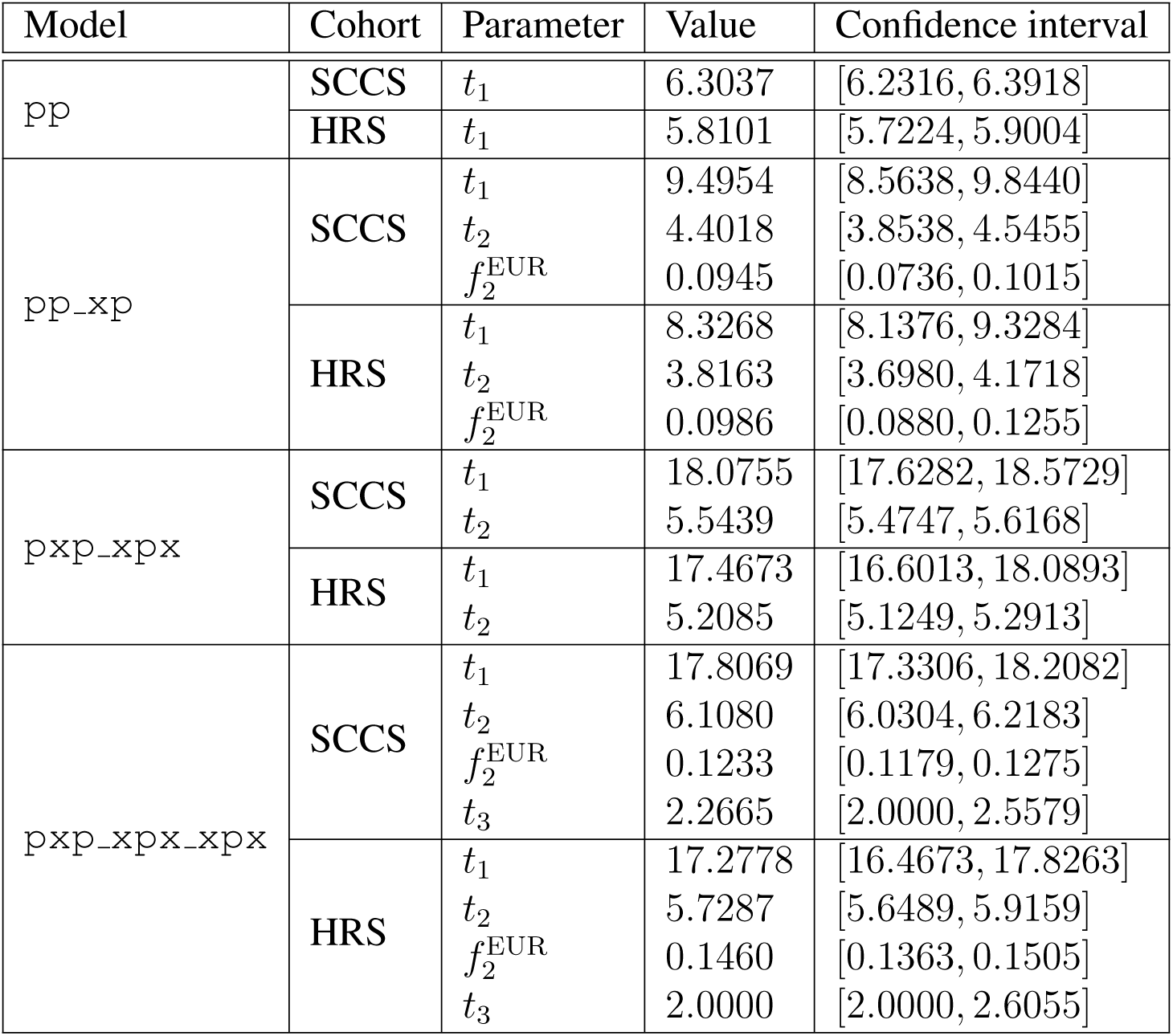
Confidence intervals for selected models inferred using Tracts. Here, *t_i_* refers to the time of the *i*th migration event (in generations ago), and 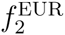 refers to the fraction of European admixture in the second migration event.

| Model | Cohort | Parameter | Value | Confidence interval |
| --- | --- | --- | --- | --- |
| pp | SCCS | $t_1$ | 6.3037 | [6.2316, 6.3918] |
| | HRS | $t_1$ | 5.8101 | [5.7224, 5.9004] |
| pp_xp | SCCS | $t_1$ | 9.4954 | [8.5638, 9.8440] |
| | | $t_2$ | 4.4018 | [3.8538, 4.5455] |
| | | $f_2^{\text{EUR}}$ | 0.0945 | [0.0736, 0.1015] |
| | HRS | $t_1$ | 8.3268 | [8.1376, 9.3284] |
| | | $t_2$ | 3.8163 | [3.6980, 4.1718] |
| | | $f_2^{\text{EUR}}$ | 0.0986 | [0.0880, 0.1255] |
| pxp_xpx | SCCS | $t_1$ | 18.0755 | [17.6282, 18.5729] |
| | | $t_2$ | 5.5439 | [5.4747, 5.6168] |
| | HRS | $t_1$ | 17.4673 | [16.6013, 18.0893] |
| | | $t_2$ | 5.2085 | [5.1249, 5.2913] |
| pxp_xpx_xpx | SCCS | $t_1$ | 17.8069 | [17.3306, 18.2082] |
| | | $t_2$ | 6.1080 | [6.0304, 6.2183] |
| | | $f_2^{\text{EUR}}$ | 0.1233 | [0.1179, 0.1275] |
| | | $t_3$ | 2.2665 | [2.0000, 2.5579] |
| | HRS | $t_1$ | 17.2778 | [16.4673, 17.8263] |
| | | $t_2$ | 5.7287 | [5.6489, 5.9159] |
| | | $f_2^{\text{EUR}}$ | 0.1460 | [0.1363, 0.1505] |
| | | $t_3$ | 2.0000 | [2.0000, 2.6055] |

**Table 4:** Characteristics of African-Americans in the HRS and SCCS cohorts.

| Cohort | African-American individuals | males / females | Hispanics | locale |
| --- | --- | --- | --- | --- |
| HRS | 1501 | 531 / 970 | 10 | contiguous US |
| SCCS | 2128 | 1131 / 997 | N/A | southern US |

**Table 5:**
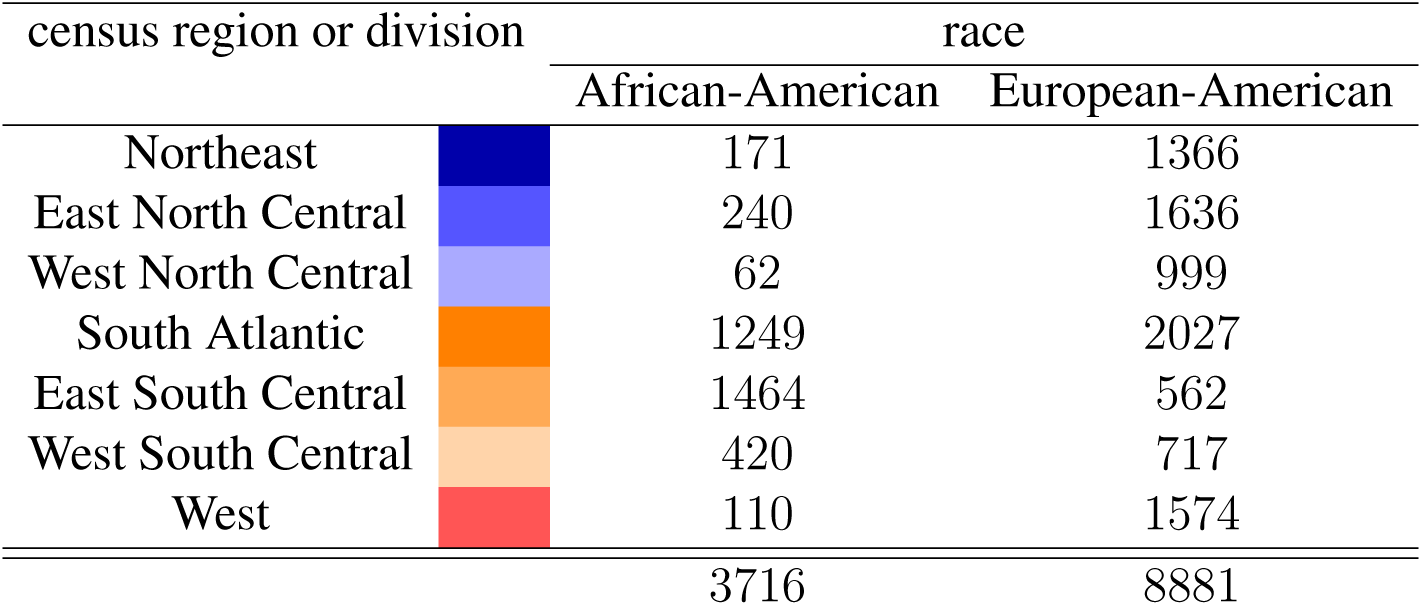
Number of US-born non-Hispanic individuals (HRS, SCCS, and ASW combined) by race and census region or division of residence in 2010 (color coded to match the maps shown in the main text).

| census region or division |  | race |  |
| --- | --- | --- | --- |
|  |  | African-American | European-American |
| Northeast |  | 171 | 1366 |
| East North Central |  | 240 | 1636 |
| West North Central |  | 62 | 999 |
| South Atlantic |  | 1249 | 2027 |
| East South Central |  | 1464 | 562 |
| West South Central |  | 420 | 717 |
| West |  | 110 | 1574 |
|  |  | 3716 | 8881 |

#### 2.3 Detailed geographic location of individuals

##### 2.3.1 HRS

The restricted HRS data contains ZIP codes for each individual, but not states. To calculate per-state global ancestry proportions within HRS, we use the following commands in Mathematica (version 10.1.0) to get the list of ZIP codes within each state in the contiguous US. For each state, we select HRS individuals whose ZIP codes belong to that state, then estimate the mean ancestry proportions for the state using the selected individuals.

~~~
states = CountryData["UnitedStates", "Regions"];
states = Delete[states, {Position[states, "Alaska"][[1]], Position[states, "Hawaii"][[1]]}];
ZIPcodes = GeoEntities[Entity["USState", #], "ZIPCode"][[All, 2]]& /@ states;
~~~

To find the spatial distance between HRS individuals, we use the ZIP Code Tabulation Area (ZCTA) database^5^ to assign latitude and longitude coordinates to the individuals based on their ZIP codes. Each coordinate in the ZCTA database is essentially the latitude and longitude coordinates of the geographic centroid of the corresponding ZIP Code Tabulation Area, as defined by the US Census Bureau. We then calculate the geodesic distance between a pair of individuals given their assigned geographic coordinates, as an estimate for the actual distance between the two individuals.

**Figure 12:**
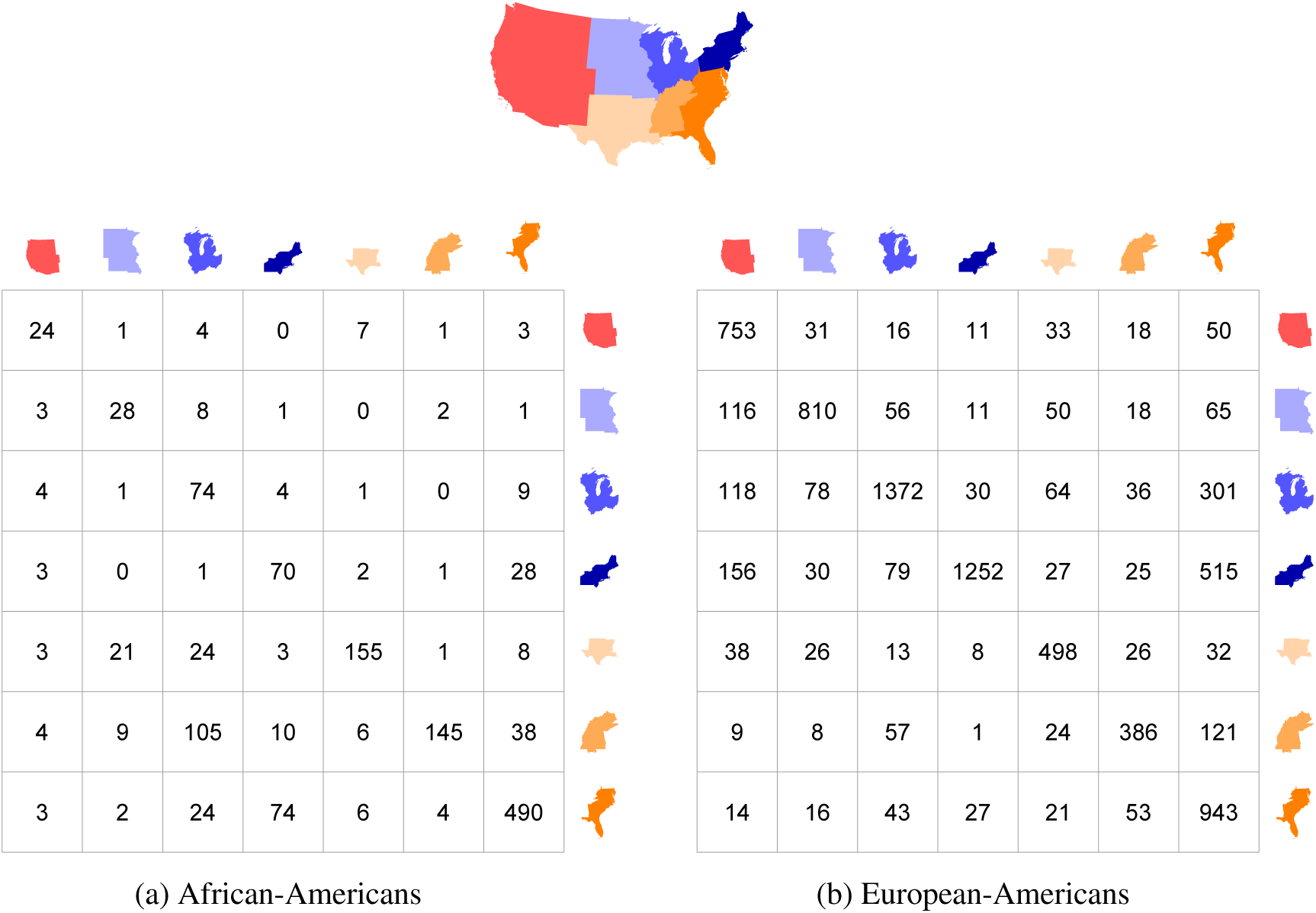
Number of non-Hispanic US-born individuals moving from one region to the other between their time of birth and the 2010 sampling year. Rows represent regions of birth, and columns represent regions of residence in 2010.

##### 2.3.2 SCCS

The data we received from SCCS contains latitude and longitude coordinates of clinics participating in the study. To convert the coordinates into specific locations (e.g., ZIP codes, states, and census regions), we used the Nominatim service from OpenStreetMap^6^ to perform reverse geocoding of the coordinates. Specifically, we used the OpenStreetMap API provided through MapQuest^7^ due to its unlimited usage policy.

#### 2.4 Principal component analysis

The results of a PCA analysis on the combined HRS, SCCS, and ASW are shown in Fig. S13.

The ASW and SCCS samples cluster with the African-American samples in HRS, as expected. The vertical axis shows that African ancestry in African-Americans varies continuously in all cohorts. African-Americans with Hispanic ethnicity are positioned slightly away from (towards the right of) the cluster of non-Hispanic African-Americans, in the same direction as other non-African-American Hispanic individuals and along the axis corresponding to Native American or Asian component. Interestingly, there are individuals in the ASW cohort with very high levels of Native American or Asian component. Specifically, 1 ASW sample lies almost halfway between the European and the Asian cluster, with almost no African component present, and 4 ASW samples have very high proportions of Native American or Asian component. Similarly, within HRS, 5 African-American samples – who have *not* self-identified as Hispanics – have very high proportions of Native American or Asian component with 4 of them having extremely low African component. Analogous to these 4 samples, there is one African-American sample – who has self-identified as Hispanic – who has a similar pattern of African, European, and Native American or Asian ancestry.

**Figure 13:**
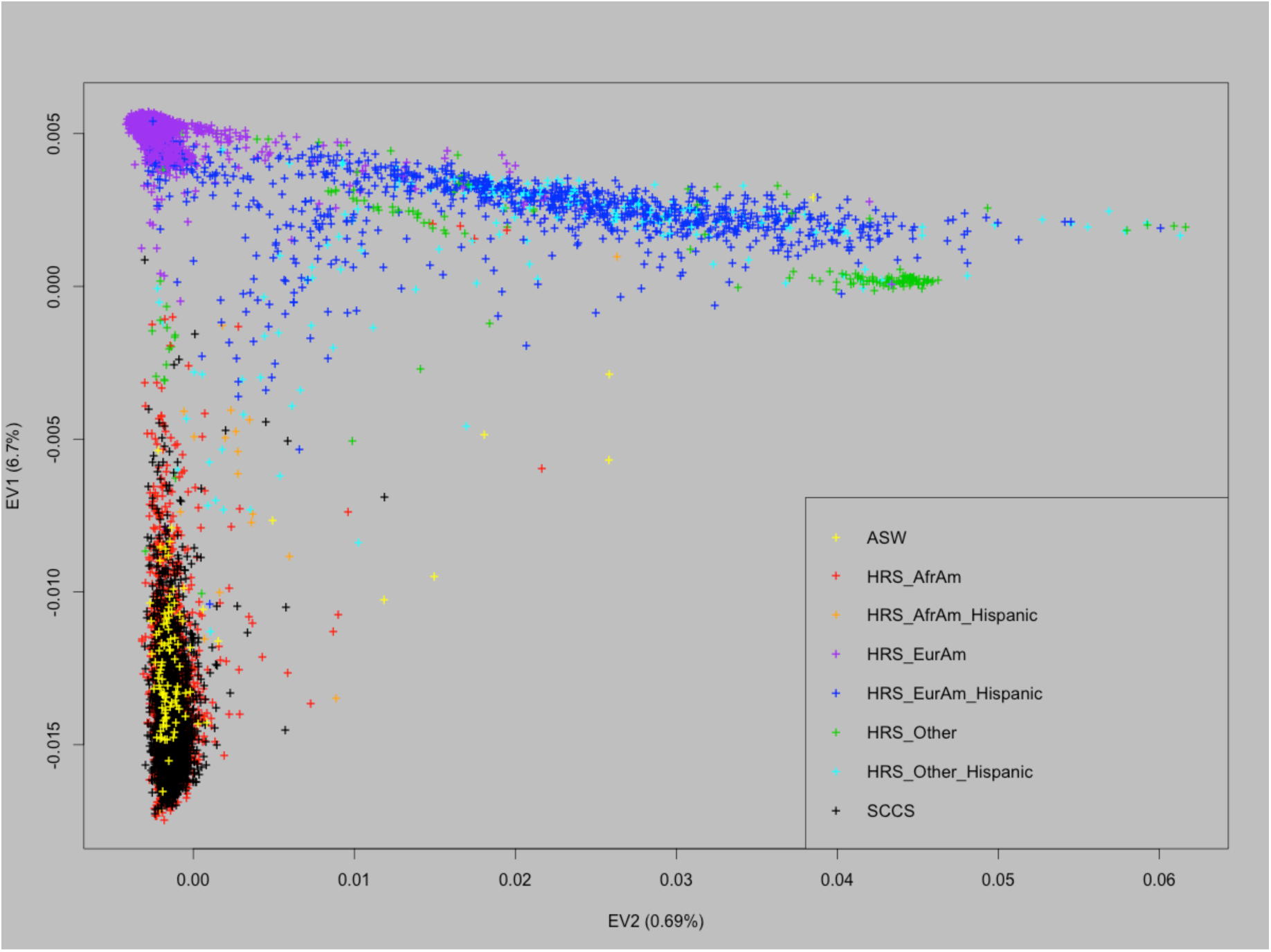
Principal component analysis of all samples in the HRS, SCCS, and ASW cohorts, using SNPRelate (*35*). The vertical axis (containing most of the variance in the data) corresponds to the distribution of African versus European component, whereas the horizontal axis indicates the distribution of Native American or Asian versus European component.

#### 2.5 Global ancestry estimates

We used the local ancestry estimates obtained from RFMix to calculate global ancestry proportions for the HRS, SCCS, and ASW cohorts as follows. For each individual, the sum of the lengths of all tracts of certain continental ancestry (i.e., African, European, or Native American) is calculated across all chromosomes from the output of RFMix and is represented as the percentage of the total length of the genome (see Fig. S14).

For the X chromosome, a supervised run of ADMIXTURE with *K* = 3 reference populations (YRI representing African ancestry, CEU representing European ancestry, and CHS representing Native American/Asian ancestry) reveals the ancestry pattern shown in Fig. S15.

#### 2.6 African-Americans of Hispanic background

We performed a supervised *K* = 4 run of ADMIXTURE (36) on African-Americans from HRS, SCCS, and also on the ASW cohort from the 1000 Genomes Project, with the YRI, CHS, GBR, IBS cohorts from the 1000 Genomes Project used as the reference populations representing African, Native American/Asian, northern European, and southern European ancestral populations. Pruning for LD was performed based on the recommendations of the authors of ADMIXTURE (PLINK arguments `--indep-pairwise 50 10 0.1`). The mean ancestry proportions for African-Americans in HRS, as estimated by ADMIXTURE, are 81.583% for African, 17.333% for European (southern and northern combined), and 1.083% for Native American, in very good agreement with those derived using local ancestry estimates of RFMix (see main text). In comparison, the ancestry proportions for the ASW cohort are found to be 75.726% for African, 21.881% for European (southern and northern combined), and 2.394% for Native American.

Fig. S16 depicts the ancestry estimates for African-Americans in the ASW, HRS, and SCCS cohorts respectively, sorted by their Native American proportions (shown in yellow). The top figure (corresponding to ASW individuals) shows that individuals with higher proportion of southern European ancestry (shown in green) tend to also have a higher proportion of Native American ancestry, and this pattern is repeated in the other two plots as well. This is especially true for HRS African-Americans who have self-identified as Hispanics (marked by the small black arrows in the middle plot), suggesting a positive correlation between the two ancestry proportions. Plotting the proportion of southern European ancestry within the total European ancestry versus the Native American ancestry for HRS African-Americans of Hispanic ethnicity reveals this correlation more clearly, as depicted by the red dots in Fig. S17. Note the presence of individuals who have *not* self-identified as Hispanics but have high proportions of both southern European and Native American ancestries. Moreover, SCCS African-Americans from Louisiana exhibit a similar pattern, as depicted by the black dots in Fig. S17.

**Figure 14:**
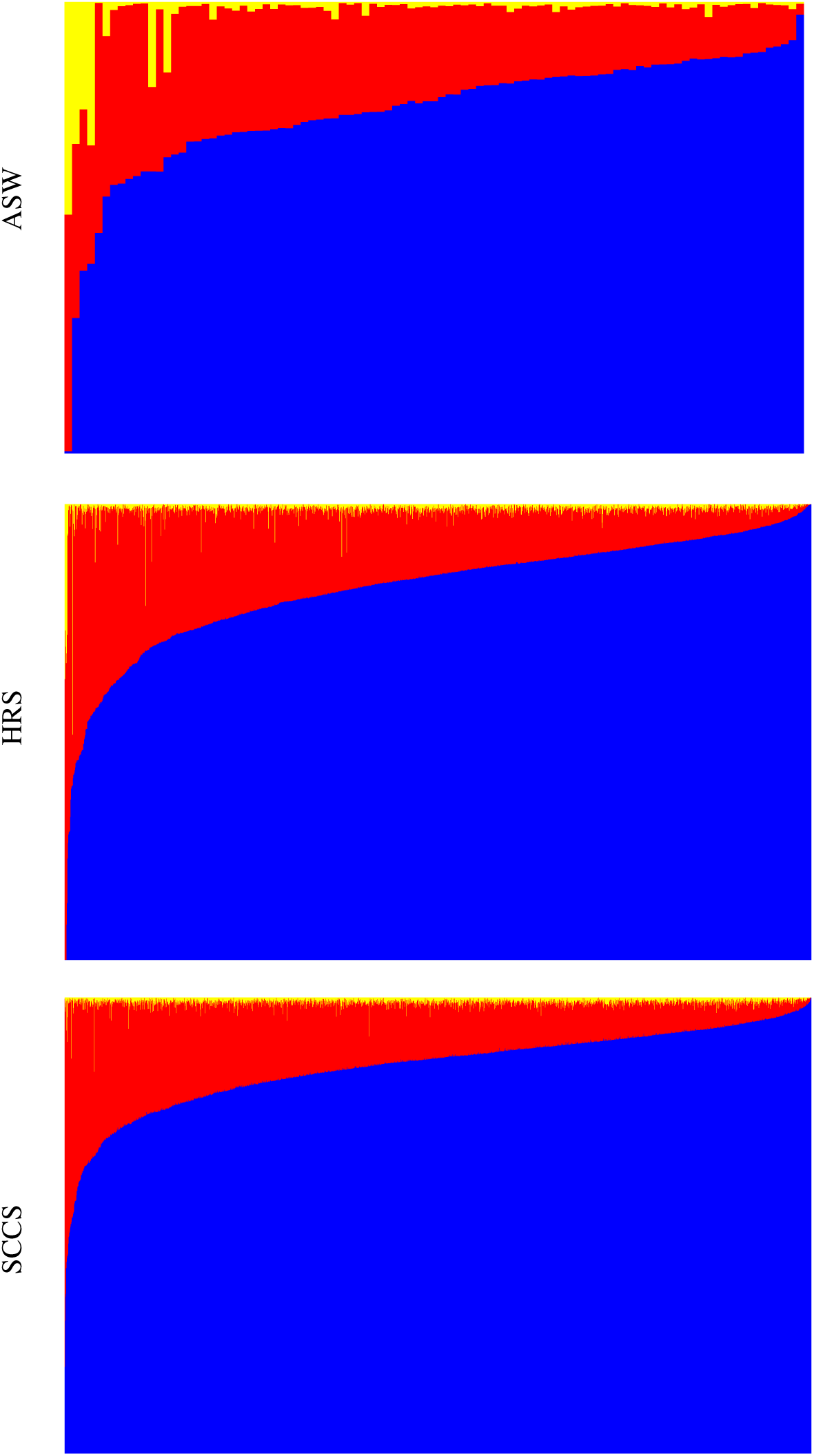
Global ancestry proportions of ASW, African-Americans in HRS, and SCCS individuals, calculated using the RFMix-inferred local ancestry. Blue, red, and yellow respectively denote African, European, and Native American or Asian ancestries. Each vertical line represent one individual, and the height of the color bars denoted the percentage of their respective ancestries in that individual.

**Figure 15:**
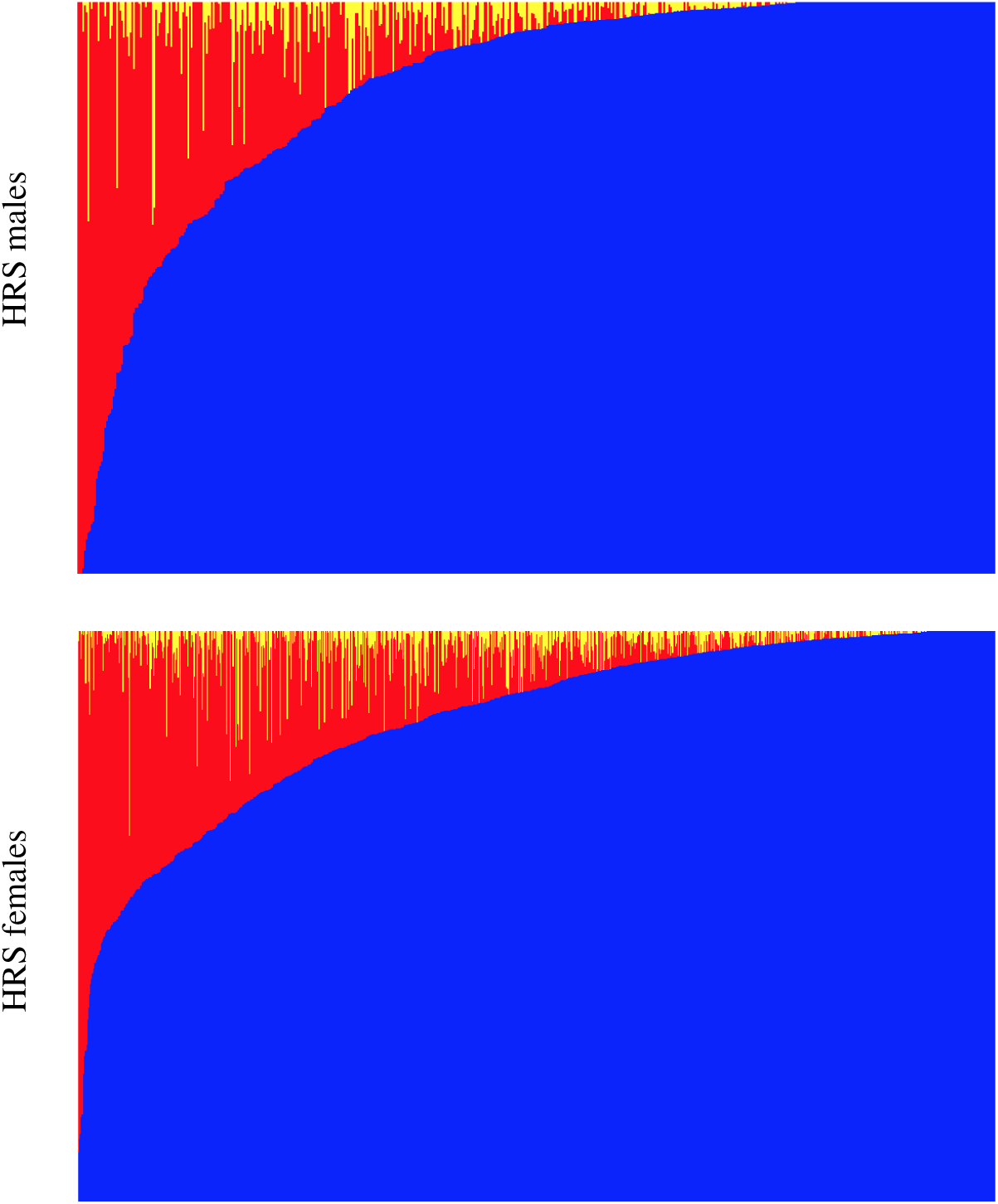
Global ancestry proportions on the X chromosome for HRS African-American males (top) and females (bottom). Each vertical bar represent one individual. Blue, red, and yellow respectively denote African, European, and Native American or Asian ancestries.

**Figure 16:**
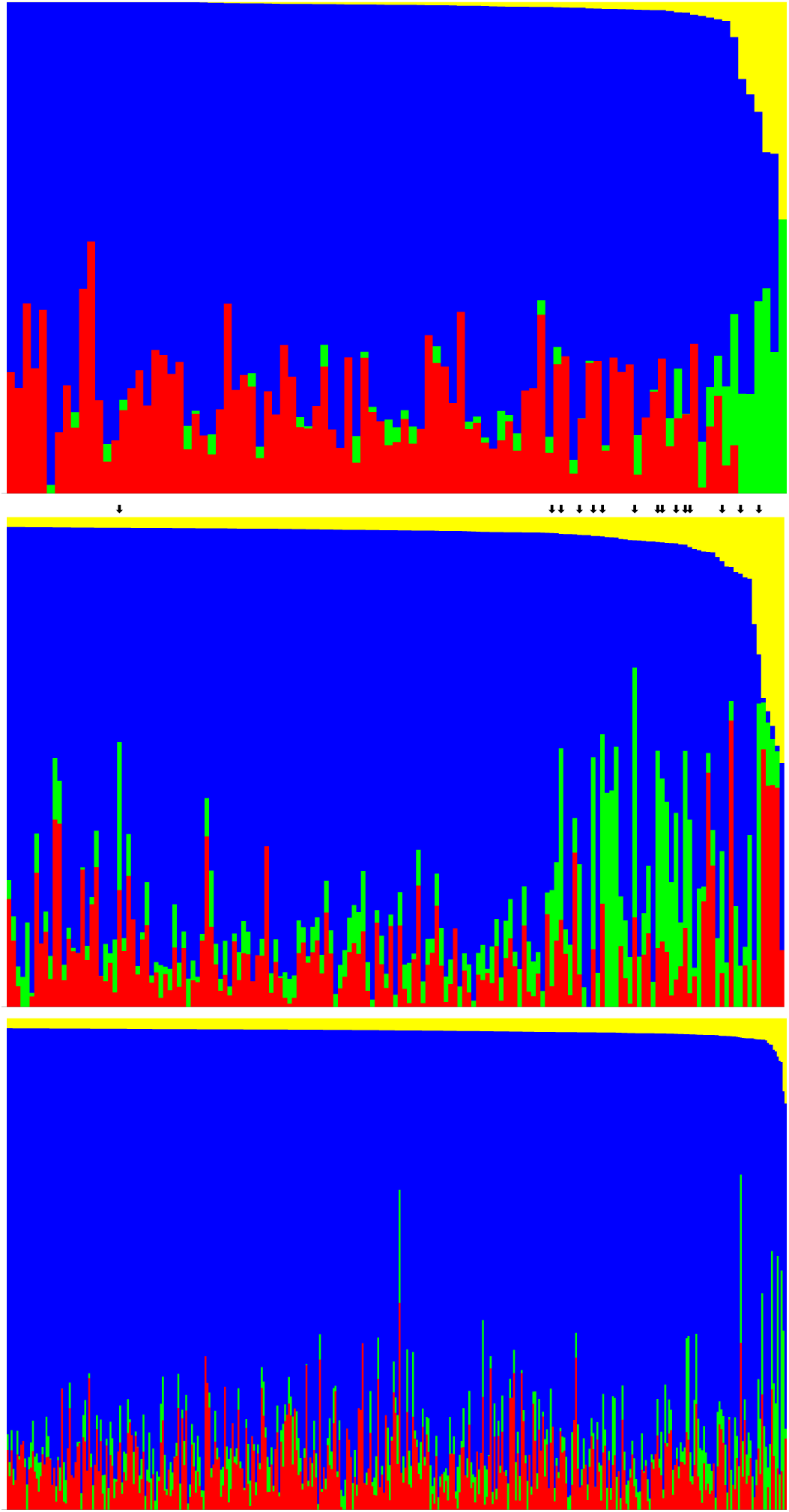
Global ancestry estimates for all ASWs (top) and for individuals with more than 2% Native American ancestry in HRS (middle) and SCCS (bottom). Yellow, blue, red, and green represent, respectively, Native American, African, northern European, and southern European ancestries. Each column represent one individual. Individuals denoted by arrows in the middle plot are self-identified Hispanic African-Americans in HRS.

**Figure 17:**
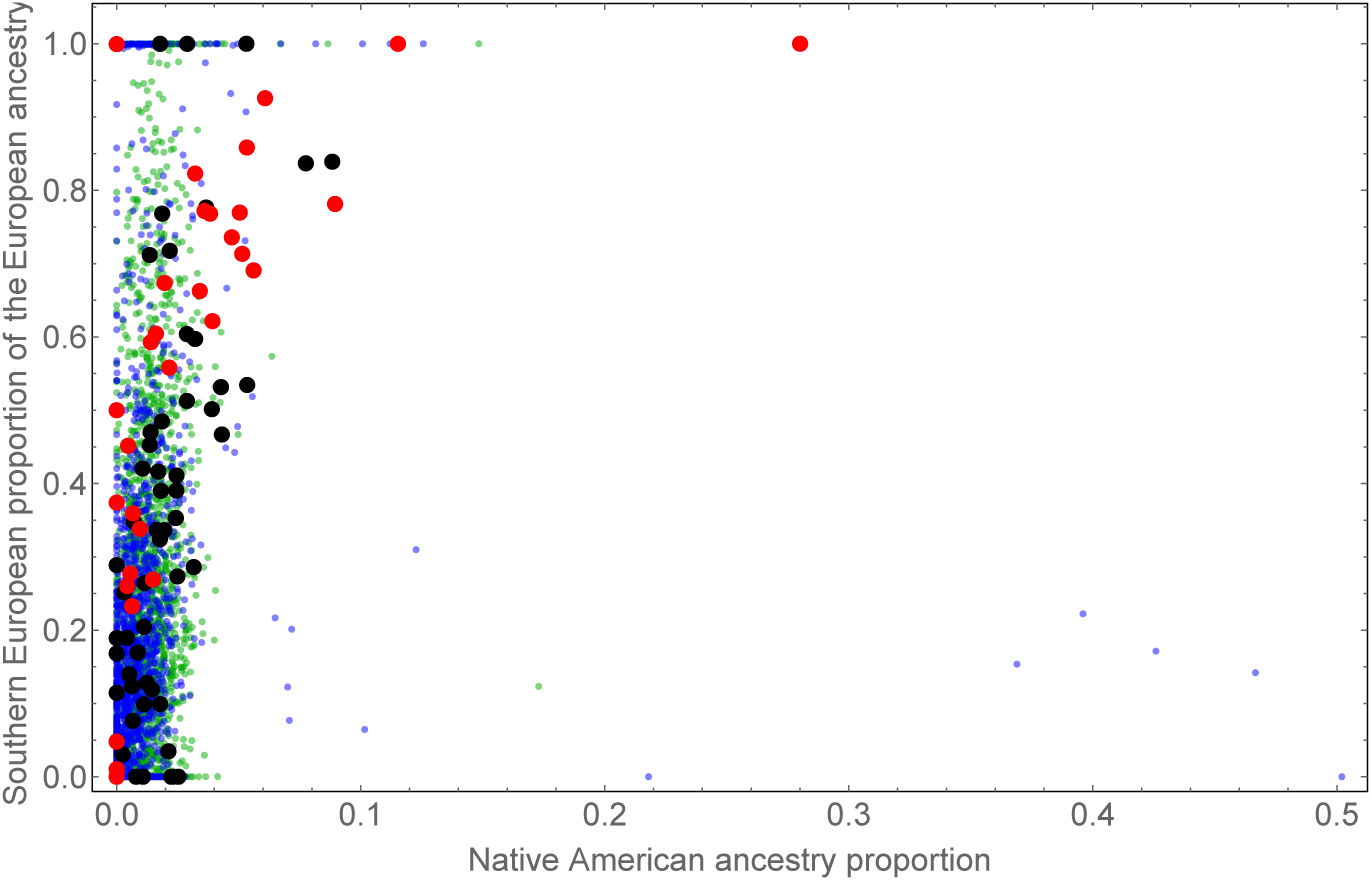
Inferred proportion of southern European ancestry within the total European ancestry versus that of Native American ancestry for African-Americans. Red represents self-identified Hispanic African-Americans in HRS, black represents SCCS African-Americans in Louisiana, and blue and green correspond, respectively, to other HRS and SCCS African-Americans.

#### 2.7 Time of admixture by region

In Fig. S18, we depict the estimated generation time since admixture in HRS for each census region, assuming a model with a single pulse of admixture.

#### 2.8 List of related individuals

We have found the pairs of individuals denoted in Table 6 to have kinship coefficients of 0.1 or greater, as estimated by PLINK. To be consistent with the definition from HRS, we have therefore labeled these pairs as related individuals and have excluded their contributions from our IBD analyses (see Methods).

**Figure 18:**
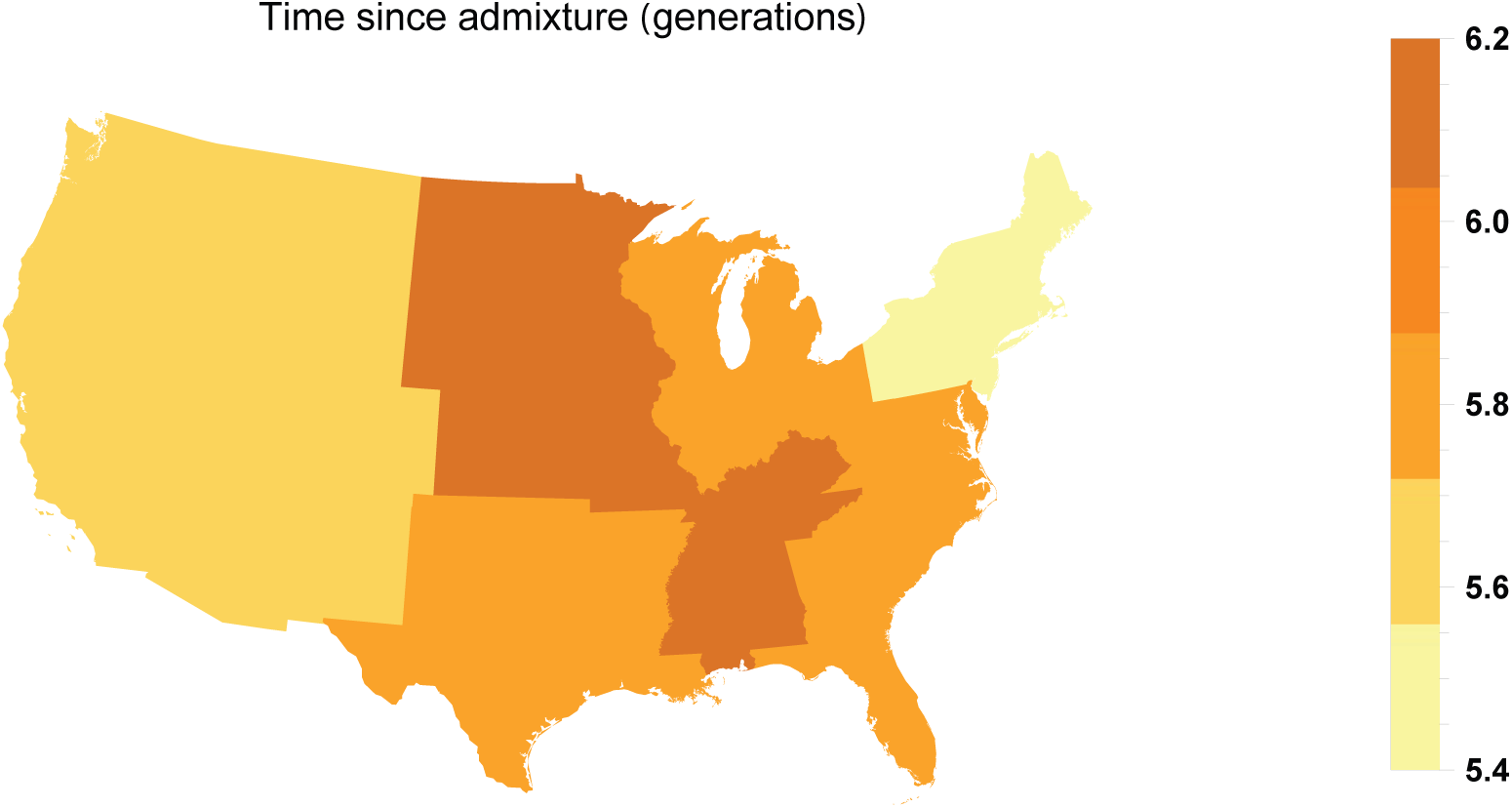
Estimated number of generation since admixture in HRS by region, assuming a single admixture pulse model in each region.

#### 2.9 Distribution of shared IBD tracts

For each individual in the HRS, SCCS, and ASW cohorts, we calculate the number of IBD segments that it shares with *all other* non-related individuals across all cohorts, using the sparse relatedness matrix 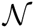 calculated above which contains the total *number* of shared IBD segments between each pair of individuals. This distribution of IBD sharing for segments in different length bins is shown in Fig. S19. For short segments, Europeans show substantially more IBD compared to African-Americans, and there is more variation in the amount of IBD. The difference is due to differences in sample size and historical effective population size. The difference in variance could reflect the greater contributions of more recent migrants in European-Americans, or more generally the presence of population structure that persisted throughout US history. By contrast, African-Americans have more long IBD, on average. Since long segments represent recent history, we attribute this difference between short and long segments to the a much greater reduction in effective population size in African-Americans compared to European-Americans since the arrival into the Americas.

**Table 6:** Related pairs of individuals in the cohorts with estimated kinship coefficient of 0.1 or larger. For relateds within HRS, we used the list provided by the Health and Retirement Study.

|  |  |  |  |  |  |
| --- | --- | --- | --- | --- | --- |
| 161321930 | GWAS_0889 | NA19625 | NA20414 | NA19982 | NA19983 |
| GWAS_0028 | GWAS_2129 | NA19700 | NA19702 | NA19983 | NA19985 |
| GWAS_0223 | GWAS_2277 | NA19701 | NA19702 | NA20126 | NA20128 |
| GWAS_0229 | GWAS_2386 | NA19703 | NA19705 | NA20127 | NA20128 |
| GWAS_0240 | GWAS_1676 | NA19704 | NA19705 | NA20276 | NA20277 |
| GWAS_0252 | GWAS_2389 | NA19707 | NA19708 | NA20278 | NA20279 |
| GWAS_0387 | GWAS_2382 | NA19713 | NA19714 | NA20279 | NA20282 |
| GWAS_0567 | GWAS_2250 | NA19713 | NA19983 | NA20279 | NA20284 |
| GWAS_0773 | GWAS_1975 | NA19713 | NA19985 | NA20282 | NA20284 |
| GWAS_0784 | GWAS_1203 | NA19714 | NA19985 | NA20282 | NA20302 |
| GWAS_0851 | GWAS_1680 | NA19818 | NA19828 | NA20282 | NA20313 |
| GWAS_0894 | GWAS_1595 | NA19819 | NA19828 | NA20287 | NA20288 |
| GWAS_0959 | GWAS_1026 | NA19834 | NA19836 | NA20289 | NA20290 |
| GWAS_1104 | GWAS_1902 | NA19835 | NA19836 | NA20289 | NA20341 |
| GWAS_1137 | GWAS_1765 | NA19900 | NA19902 | NA20290 | NA20341 |
| GWAS_1168 | GWAS_1803 | NA19901 | NA19902 | NA20291 | NA20292 |
| GWAS_1323 | GWAS_1928 | NA19908 | NA19919 | NA20294 | NA20295 |
| GWAS_1375 | GWAS_2257 | NA19909 | NA19919 | NA20296 | NA20297 |
| GWAS_1380 | GWAS_1833 | NA19914 | NA19915 | NA20299 | NA20300 |
| GWAS_1571 | GWAS_1598 | NA19916 | NA19918 | NA20302 | NA20313 |
| GWAS_1596 | GWAS_2171 | NA19917 | NA19918 | NA20314 | NA20316 |
| GWAS_1962 | GWAS_2008 | NA19920 | NA20129 | NA20317 | NA20319 |
| GWAS_2015 | GWAS_2226 | NA19921 | NA20129 | NA20332 | NA20333 |
| NA20332 | NA20343 | NA20342 | NA20343 | NA20357 | NA20358 |
| NA20334 | NA20335 | NA20344 | NA20345 | NA20359 | NA20360 |
| NA20334 | NA20336 | NA20344 | NA20350 | NA20359 | NA20363 |
| NA20334 | NA20337 | NA20346 | NA20347 | NA20363 | NA20364 |
| NA20335 | NA20336 | NA20347 | NA20363 |  |  |
| NA20336 | NA20337 | NA20356 | NA20358 |  |  |

**Figure 19:**
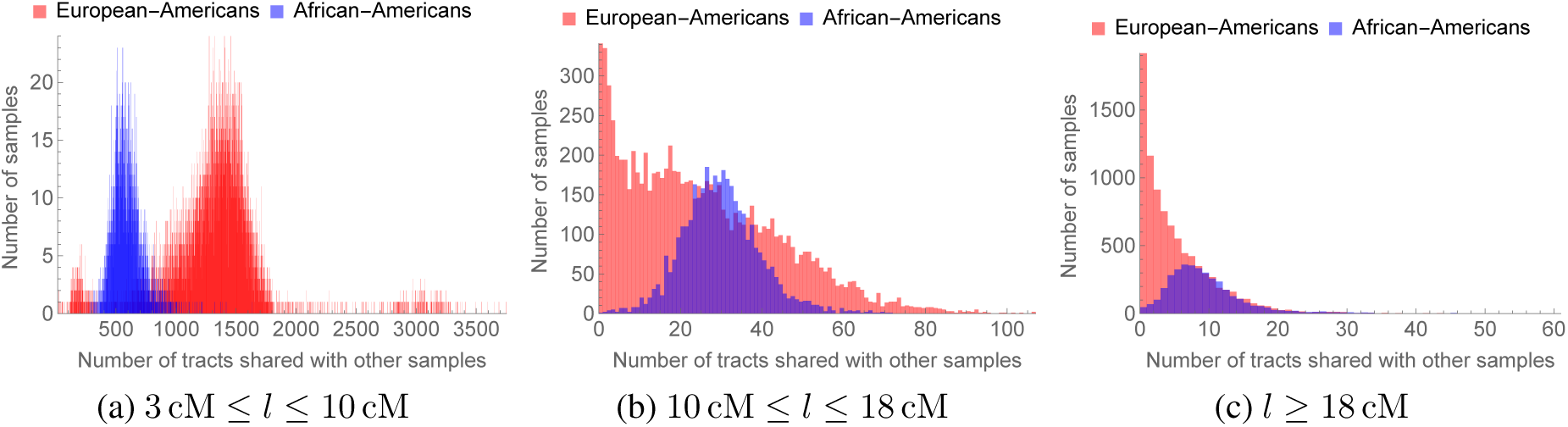
Distribution of IBD sharing for African-American (blue) and European-American (red) individuals using IBD tracts belonging to different length bins.

IBD sharing for European-Americans tends to be mostly via shorter (and, therefore, older) IBD segments compared to that for African-Americans. Hence, for bins containing longer IBD segments, the peak corresponding to European-American IBD sharing moves to the left faster compared to the respective peak for African-Americans.

#### 2.10 Regional IBD relatedness and sampling locations

Using the individuals’ region of residence in 2010 as their location, we find the relatedness pattern shown in Fig. S20 between census regions for African-Americans and European-Americans. On the other hand, using the individuals’ region of birth as their location instead of their 2010 region of residence, we find the relatedness pattern shown in Fig. S21 between census regions for African-Americans and European-Americans.

The differences between Fig. S21 and Fig. S20 are due to the following factors: If we use the 2010 region of residence as opposed to the region of birth as the individuals’ locations, the number of individuals in the northern and western census regions increases due to the migrations from the South. This is the reason that more North-to-West connections are shown in Fig. S20 than in Fig. S21, considering our criterion of a minimum of 10,000 potential IBD pairs for visualization of the relatedness between two regions. Moreover, we also see that the connection between West South Central and West North Central is weaker Fig. S20 than in Fig. S21. This is mainly due to 21 African-American individuals in HRS who were born in West South Central and who have large IBD with other individuals who were born in northern regions, especially those in West North Central. These individuals later moved to West North Central and, thus, are sampled in the latter region in 2010 (see Fig. S12a). The observed depletion of the IBD signal connecting the aforementioned two regions (which is explained by these migrations) is, thus, a sampling effect.

**Figure 20:**
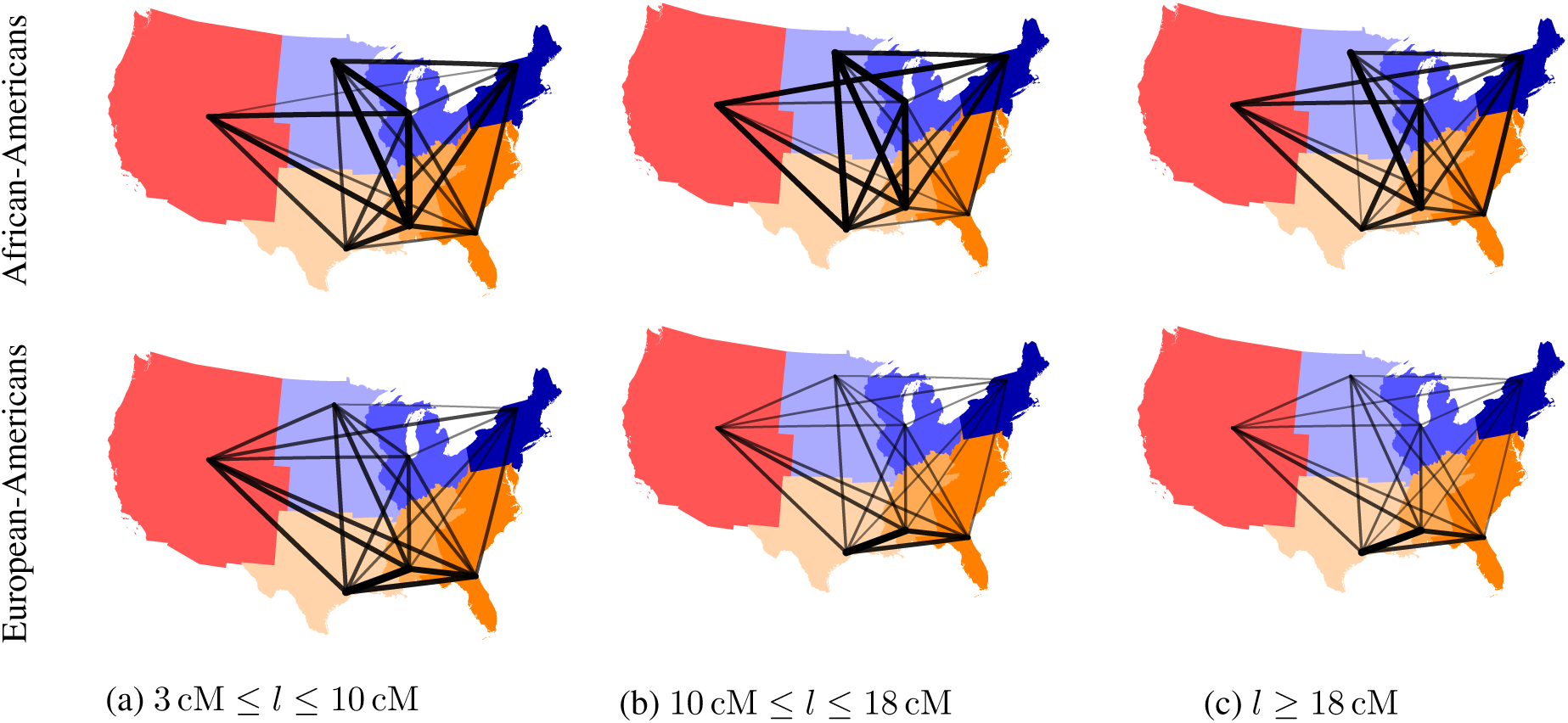
IBD relatedness among African-Americans (top row) and among European-Americans (bottom row) across the US census regions (using 2010 region of residence). In each subfigure, the thickness and opacity of the line connecting any two regions show the strength of relatedness between those regions. Note that scaling of lines is not equal across different subfigures, and relatedness between regions with fewer than 10,000 possible pairs of individuals is not shown (see Methods for details).

**Figure 21:**
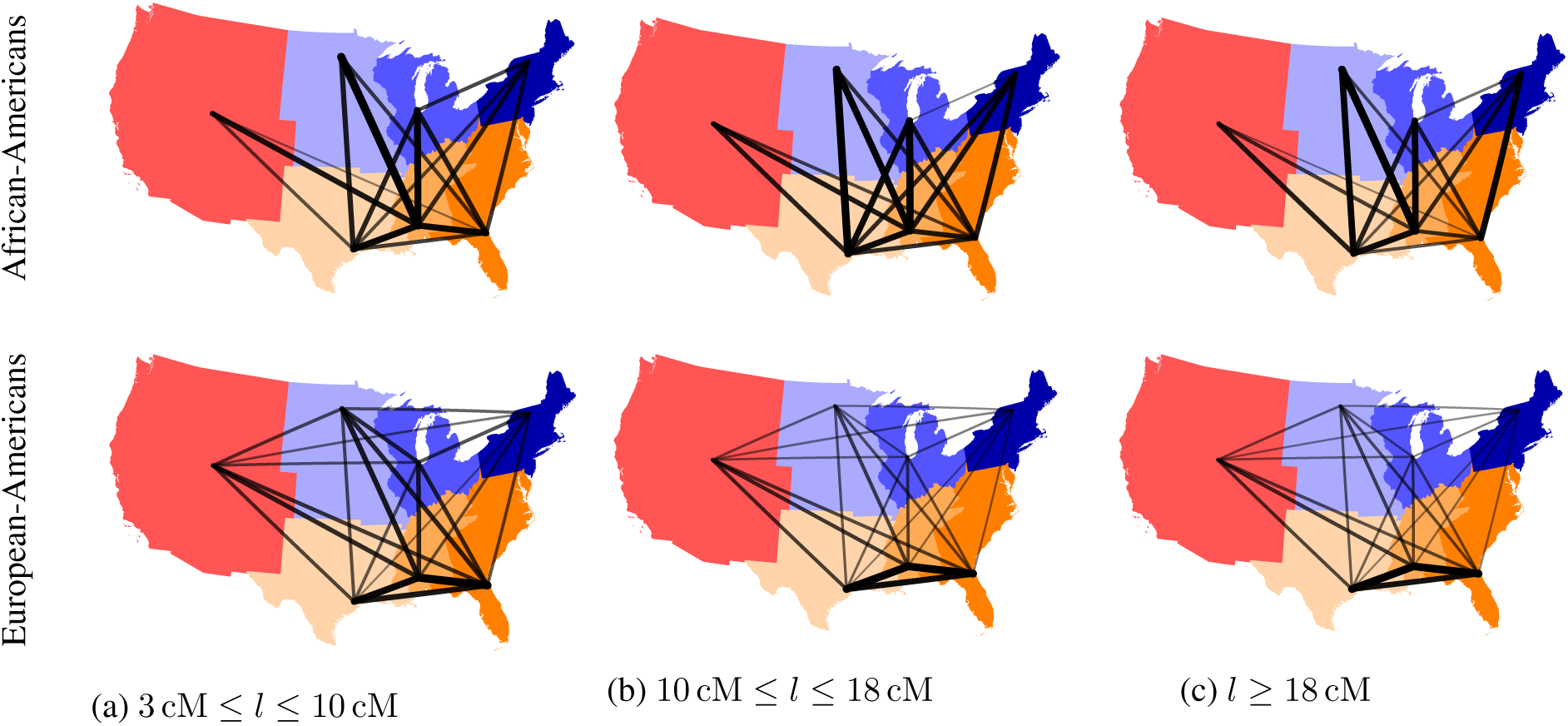
IBD relatedness among African-Americans (top row) and among European-Americans (bottom row) across the US census regions (using the regions of birth). In each subfigure, the thickness and opacity of the line connecting any two regions show the strength of relatedness between those regions. Note that scaling of lines is not equal across different subfigures, and relatedness between regions with fewer than 10,000 possible pairs of individuals is not shown (see Methods for details).

Fig. S22 shows details of the relatedness patterns among African-Americans across US census regions. The grayscale plots in the top row show the average pairwise IBD length shared between census regions, calculated using all IBD segments satisfying the length criteria shown below each column. Actual values are shown in the middle row, whereas plots in the bottom row show the total number of IBD segments shared between the US census regions. Relatedness between European-Americans across US census regions is similarly displayed in Fig. S23. Relatedness between African-Americans and European-Americans is shown in Fig. S24.

#### 2.11 Census-based regional relatedness

Census-based relatedness between the US regions, estimated via the 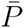 (directional) and *I* (non-directional) metrics (as defined in the Methods), are shown in Fig. S25 for African-Americans and European-Americans.

**Figure 22:**
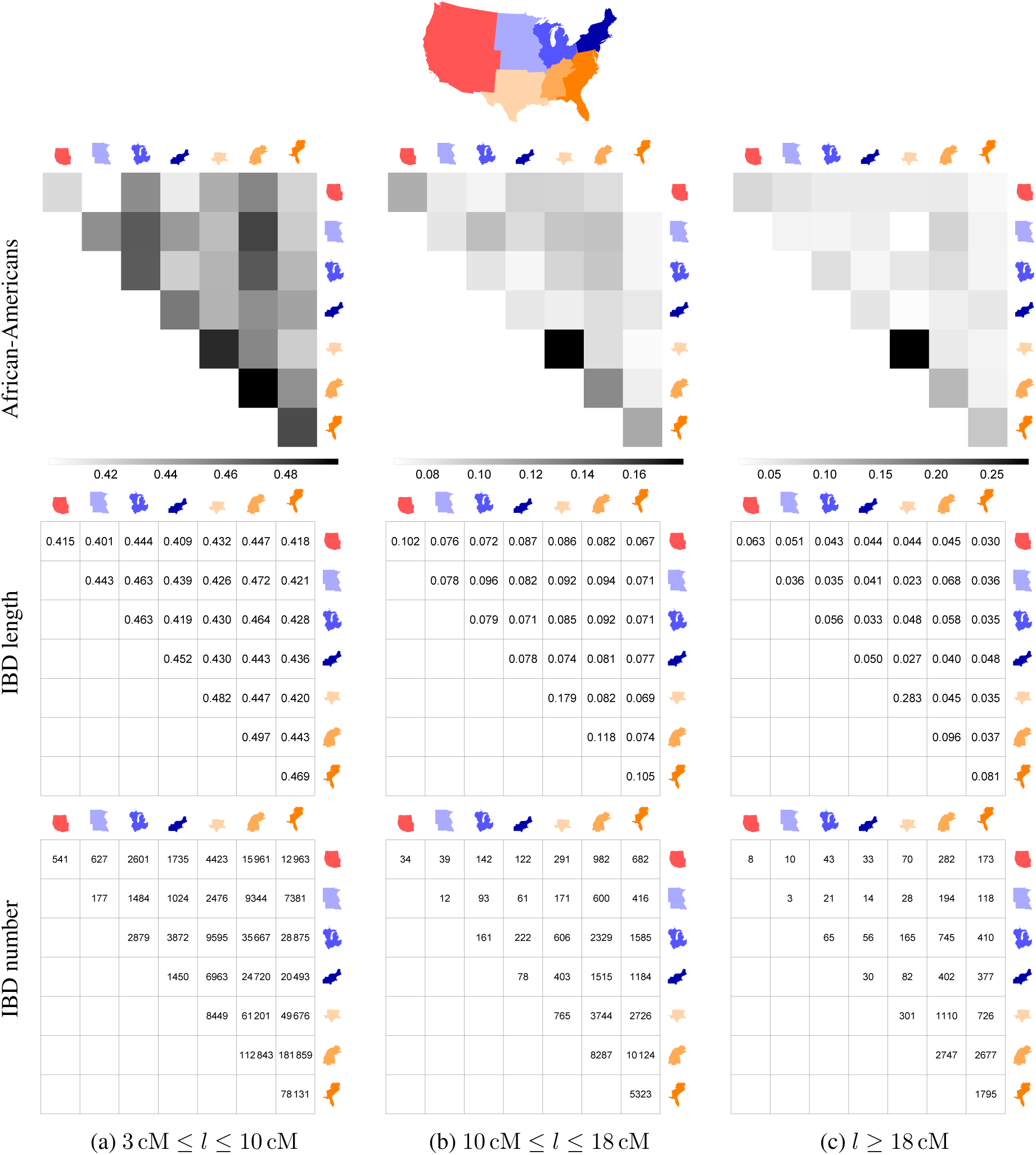
Relatedness between African-Americans across US census regions based on the average total length of shared IDB segments of length in the specified ranges (using region of residence in 2010). The values shown in the second row are converted to grayscale in the top row to aid visualization, with the scales presented underneath each figure. Since the matrices are symmetric, only the upper-triangular parts are shown.

**Figure 23:**
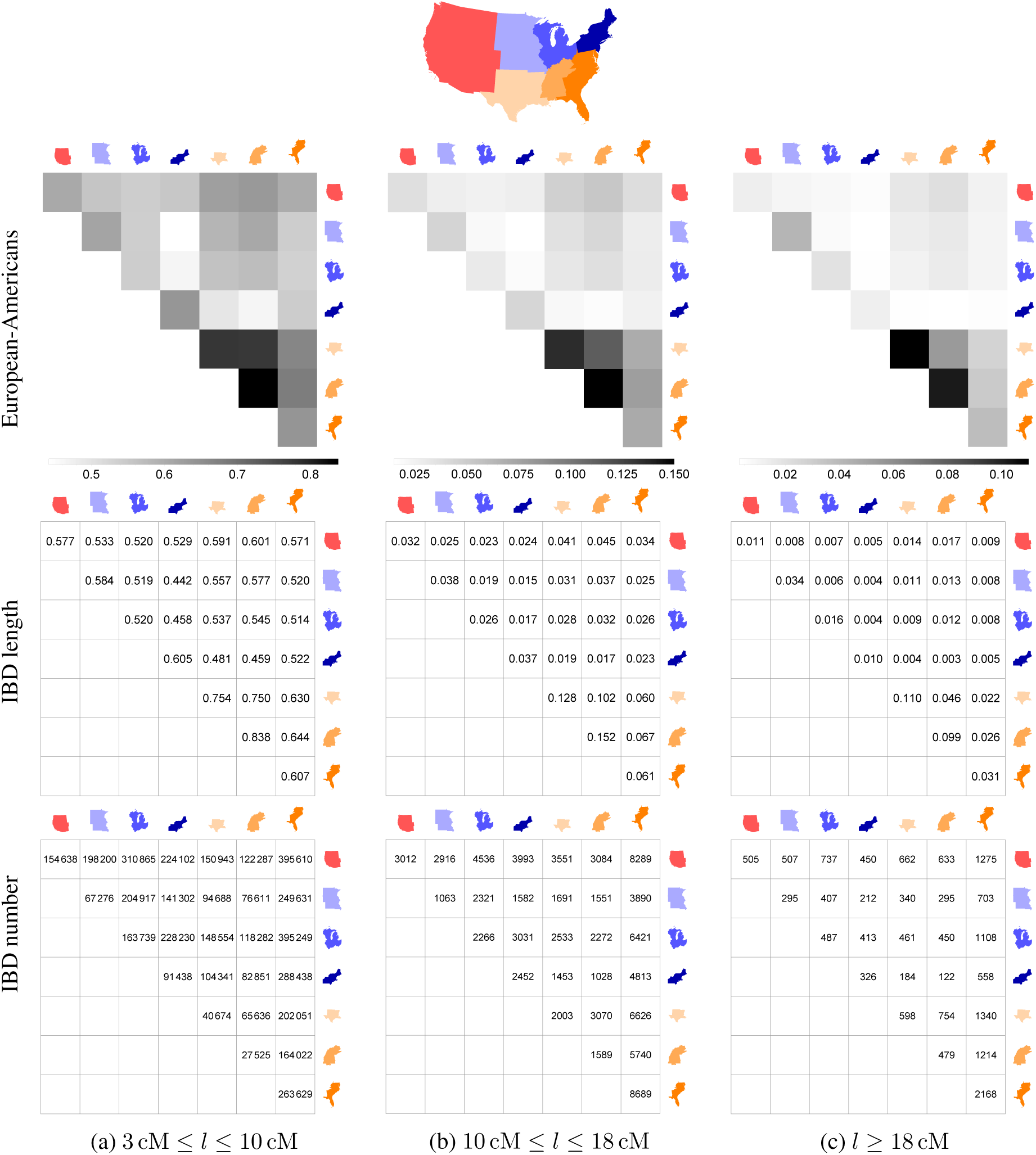
Relatedness between European-Americans across US census regions based on the average total length of shared IDB segments of length in the specified ranges (using region of residence in 2010). The values shown in the second row are converted to grayscale in the top row to aid visualization, with the scales presented underneath each figure. Since the matrices are symmetric, only the upper-triangular parts are shown.

**Figure 24:**
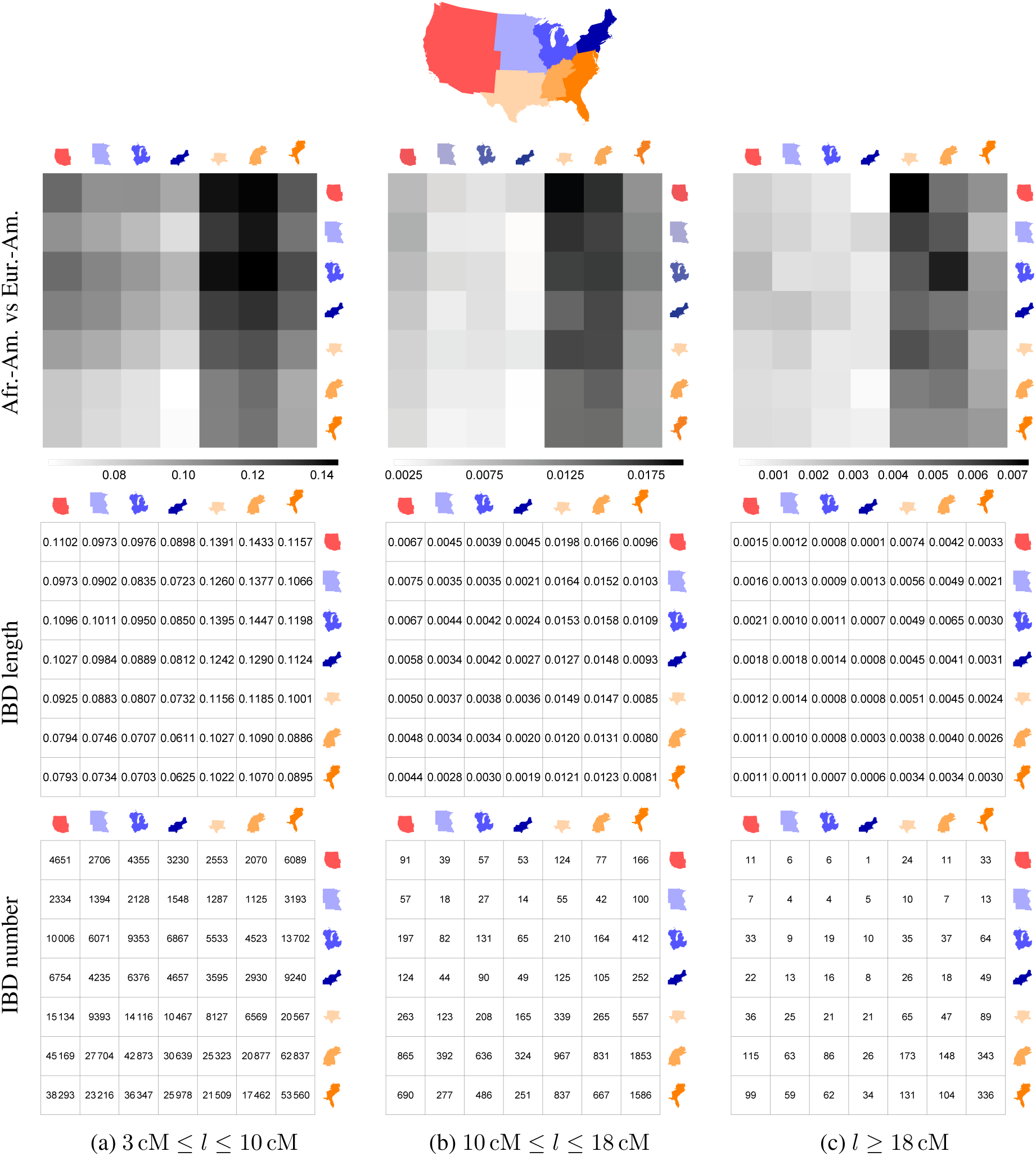
Relatedness between African-Americans and European-Americans across US census regions based on the average total length (top and middle rows) and number (bottom row) for IDB segments of length in the specified ranges (using region of residence in 2010). The values shown in the second row are converted to grayscale in the top row to aid visualization, with the scales presented underneath each figure. The columns in each figure represent European-Americans, and the rows represent African-Americans.

**Figure 25:**
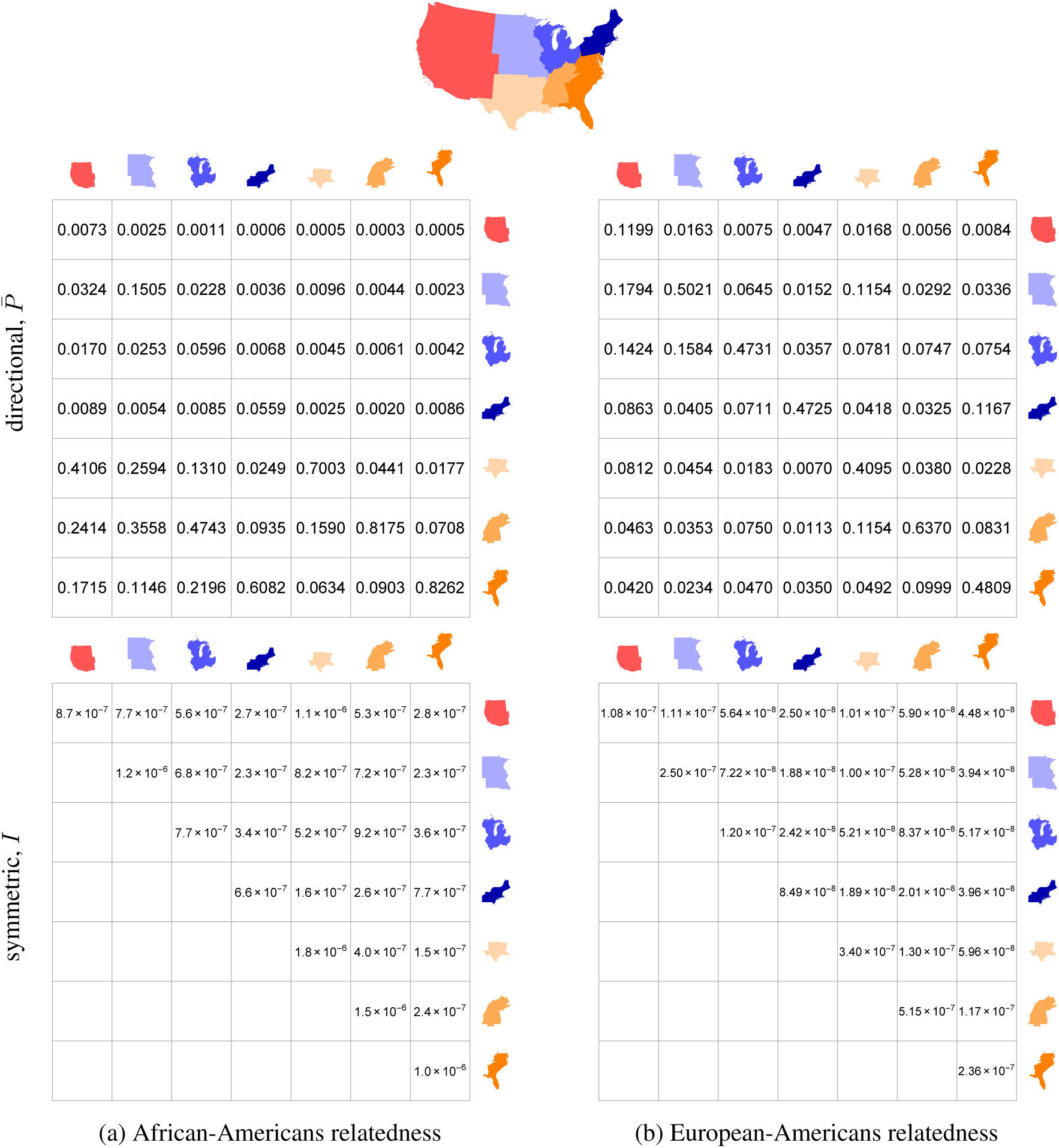
Census-based predicted relatedness between (a) African-Americans and (b) European-Americans across the US census regions. The top row shows the values for the directional metric 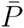, whereas the bottom row shows those for the symmetric one *I*. In the top figures (read column-wise), each column shows for its respective census region the proportion of ancestral population which originated from other census regions.

## Footnotes

1 ftp://ftp.1000genomes.ebi.ac.uk/vol1/ftp/release/20130502/supporting/hd_genotype_chip/

2 http://mathgen.stats.ox.ac.uk/impute/data_download_1000G_phase1_integrated_SHAPEIT2_9–12–13.html

3 For connection between a random walk and the diffusion model, see, e.g., http://ocw.mit.edu/courses/mathematics/18–366-random-walks-and-diffusion-fall-2006/index.htm

4 ftp.1000genomes.ebi.ac.uk/vol1/ftp/phase1/analysis_results/supporting/omni_haplotypes/

5 From the 2014 US Gazetteer Files by the US Census Bureau https://www.census.gov/geo/maps-data/data/gazetteer2014.html

6 Data available under the Open Database Licence; © OpenStreetMap contributors.

7 Details at http://open.mapquestapi.com/nominatim/#reverse

## References

1. Voyages Database. Voyages: The Trans-Atlantic slave trade database (2009). URL http://www.slavevoyages.org.

2. Ruggles, S. et al. Integrated Public Use Microdata Series: Version 5.0 [machine-readable database]. Tech. Rep., University of Minnesota, Minneapolis (2010).

3. Wilkerson, I. The warmth of other suns: The epic story of America’s great migration (Vintage, 2010).

4. Lemann, N. The Promised Land: The Great Black Migration and How It Changed America (Vintage, 1992).

5. Bustamante, C. D., De La Vega, F. M. & Burchard, E. G. Genomics for the world. Nature 475, 163–165 (2011).

6. Burchard, E. G. Missing patients. Nature 513, 301–302 (2014).

7. Tishkoff, S. A. et al. The genetic structure and history of Africans and African Americans. Science 324, 1035–1044 (2009).

8. Parra, E. J. et al. Estimating African American admixture proportions by use of population-specific alleles. The American Journal of Human Genetics 63, 1839–1851 (1998).

9. Kidd, J. M. et al. Population genetic inference from personal genome data: Impact of ancestry and admixture on human genomic variation. The American Journal of Human Genetics 91, 660–671 (2012).

10. Bryc, K. et al. Genome-wide patterns of population structure and admixture in West Africans and African Americans. Proceedings of the National Academy of Sciences 107, 786–791 (2012).

11. Maples, B. K., Gravel, S., Kenny, E. E. & Bustamante, C. D. RFMix: A discriminative modeling approach for rapid and robust local-ancestry inference. The American Journal of Human Genetics 93, 278–288 (2013).

12. Bryc, K., Durand, E. Y., Macpherson, J. M., Reich, D. & Mountain, J. L. The genetic ancestry of African Americans, Latinos, and European Americans across the United States. The American Journal of Human Genetics 96, 37–53 (2015).

13. Zakharia, F. et al. Characterizing the admixed African ancestry of African Americans. Genome Biology 10, R141 (2009).

14. Auton, A. et al. Global distribution of genomic diversity underscores rich complex history of continental human populations. Genome Research 19, 795–803 (2009).

15. Smith, M. W. et al. A high-density admixture map for disease gene discovery in African Americans. The American Journal of Human Genetics 74, 1001–1013 (2004).

16. Gravel, S. Population genetics models of local ancestry. Genetics 191, 607–619 (2012).

17. Lind, J. M. et al. Elevated male European and female African contributions to the genomes of African American individuals. Human Genetics 120, 713–722 (2007).

18. Berlin, I. The Making of African America: The Four Great Migrations (Penguin, 2010).

19. Hayes, K. H. Slavery Before Race: Europeans, Africans, and Indians at Long Island’s Sylvester Manor Plantation (New York University Press, 2013).

20. Moreno-Estrada, A. et al. The genetics of Mexico recapitulates Native American substructure and affects biomedical traits. Science 344, 1280–1285 (2014).

21. Fejerman, L. et al. Genome-wide association study of breast cancer in Latinas identifies novel protective variants on 6q25. Nature Communications 5, 5260 (2014).

22. Ramachandran, S. et al. Support from the relationship of genetic and geographic distance in human populations for a serial founder effect originating in Africa. Proceedings of the National Academy of Sciences 102, 15942–15947 (2005).

23. The Genome of the Netherlands Consortium. Whole-genome sequence variation, population structure and demographic history of the Dutch population. Nature Genetics 46, 818–825 (2014).

24. Palamara, P. F., Lencz, T., Darvasi, A. & Pe’er, I. Length distributions of identity by descent reveal fine-scale demographic history. The American Journal of Human Genetics 91, 809–822 (2012).

25. Browning, S. R. & Thompson, E. A. Detecting rare variant associations by identity-by-descent mapping in case-control studies. Genetics 190, 1521–1531 (2012).

26. Juster, F. T. & Suzman, R. An overview of the Health and Retirement Study. The Journal of Human Resources 30, S7–S56 (1995).

27. Signorello, L. B. et al. Southern community cohort study: establishing a cohort to investigate health disparities. Journal of the National Medical Association 97, 972–979 (2005).

28. The 1000 Genomes Project Consortium. An integrated map of genetic variation from 1,092 human genomes. Nature 491, 56–65 (2012).

29. Chang, C. C. et al. Second-generation PLINK: Rising to the challenge of larger and richer datasets. GigaScience 4, 7 (2015).

30. Turner, S. et al. Quality control procedures for genome-wide association studies. Current Protocols in Human Genetics 68, 1.19.1–1.19.18 (2011).

31. Delaneau, O., Zagury, J.-F. & Marchini, J. Improved whole-chromosome phasing for disease and population genetic studies. Nature Methods 10, 5–6 (2012).

32. Gusev, A. et al. Whole population, genome-wide mapping of hidden relatedness. Genome Research 19, 318–326 (2009).

33. Durand, E. Y., Eriksson, N. & McLean, C. Y. Reducing pervasive false-positive identical-by-descent segments detected by large-scale pedigree analysis. Molecular Biology and Evolution 31, 2212–2222 (2014).

34. Arfken, G. B. & Weber, H. J. Mathematical Methods for Physicists (Academic Press, Orlando, FL, 1985), 3rd edn.

35. Zheng, X. et al. A high-performance computing toolset for relatedness and principal component analysis of SNP data. Bioinformatics 28, 3326–3328 (2012).

36. Alexander, D. H., Novembre, J. & Lange, K. Fast model-based estimation of ancestry in unrelated individuals. Genome Research 19, 1655–1664 (2009).

